# Dual mechanism of kinetochore microtubule detachment

**DOI:** 10.1101/2023.06.05.543640

**Authors:** William Conway, Gloria Ha, Luyi Qiu, Ariel Amir, Daniel J. Needleman

## Abstract

During eukaryotic cell division, microtubules connect to chromosomes by attaching to the kinetochore via the NDC80 complex (NDC80c). Improperly attached kinetochore microtubules (KMTs) are selective detached from the kinetochore. While the proper regulation of KMT detachments is crucial for accurate chromosome segregation, the manner by which KMT detachments are controlled by the biophysical properties of NDC80 and KMTs remains poorly understood. Here, we investigate the mechanism of KMT detachment by combining quantitative measurements of NCD80c-KMT binding and KMT detachments, with FLIM-FRET and photoconversion, and mathematical modeling. We find that individual NDC80c at kinetochores bind noncooperatively to KMTs, with a free energy difference between bound and unbound NDC80c of −1.0 ± 0.1k_B_T, and we determine that each phosphorylated residue of NDC80c decreases the NDC80c-KMT binding energy by 0.35 ± 0.03k_B_T. We show that the energetics of NDC80c binding can be related to KMT detachment by a dual detachment mechanism in which KMTs detach from kinetochores when either 1) all NDC80c spontaneously unbind from the KMT or 2) following KMT catastrophe. We find that the affinity of NDC80c for KMTs is reduced at low-tension, non-bioriented kinetochores due to centromere-localized Aurora B phosphorylating the NDC80c, resulting in an elevated detachment rate for the associated KMTs. Taken together, this work leads to a multiscale, bottom-up biophysical model for how the energetics of NDC80c-KMT interactions impact KMT detachments and provides an understanding of the molecular basis of their regulation during mitotic error correction.

**SIGNIFICANCE STATEMENT:** Chromosome segregation errors can lead to aneuploidy, birth defects and cancer. Accurate cellular division, where one copy of each chromosome is segregated to each daughter cell, depends on regulation of attachments between chromosomes and microtubules in the mitotic spindle. While extensive work has identified proteins involved in suppressing mitotic errors, it is still unclear how incorrect chromosome-microtubule attachments which result in mitotic errors are selectively removed. Here, we employed a combination of quantitative live-cell fluorescence imaging, molecular perturbations, and mathematical modeling to elucidate the biophysical mechanism of how microtubules are selectively detached from misaligned mitotic chromosomes. This work leads to quantitative understanding of how chromosome attachments are regulated and mitotic errors are corrected.

## INTRODUCTION

Biological systems are organized and regulated across length scales, ranging from molecules to cells to tissues, where individual components hierarchically assemble into larger, functional structures (1–5). During cell division, molecular-scale interactions between microtubule filaments, molecular motors and chromosomes organize a cellular-scale mitotic spindle that is responsible for segregating one copy of each chromosome to each daughter cell, which in turn ensures robust development of tissues and organisms (6–10). How the regulation of individual spindle components ensures accurate chromosome segregation has remined unclear due to a lack of quantitative experiments connecting molecular inputs to cellular scale outputs that would enable the development of multiscale, bottom-up biophysical mechanisms in cell biology (3, 11–13).

The mitotic spindle is a bipolar structure composed of thousands of microtubules interacting with chromosomes and molecular motors which is responsible for segregating chromosomes during eukaryotic cell division (7–10, 14). In the spindle, microtubules bind to chromosomes at a macromolecular complex called the kinetochore and biorient the chromosomes along the spindle axis to ensure that each daughter cells receives a single copy of each chromosome (15–19). Erroneous attachments by kinetochore microtubules (KMTs) to non-bioriented chromosomes are thought to cause errors in chromosomes segregation (20–22). To ensure accurate chromosome segregation, potential errors are actively corrected during metaphase. Extensive evidence suggests that this mitotic error correction process is based on selectively detaching erroneously attached KMTs from non-bioriented kinetochores (23–25).

While there has been extensive work identifying the molecular players involved in regulating KMT attachments, the mechanism of how KMTs detach from the kinetochore is not well understood. The NDC80c, an outer kinetochore subunit, is thought to be the primary coupler between KMTs and the kinetochore (26–28).There are approximately 200 NDC80c (29) and 10 KMTs (9) at each kinetochore in human cells, so each KMT can interact with roughly 20 NDC80c. The NDC80c has been implicated as an important regulator of KMT stability, as have microtubule-depolymerizers such as MCAK, a kinesin-13 that localizes to centromeres during mitosis (15, 30–37). The relative contribution of NDC80c binding and depolymerizing kinesins in regulating KMT attachments is unclear (34, 38, 39).

Selective detachment of KMTs from erroneously attached chromosomes is thought to be crucial for mitotic error correction and is mediated by inter-kinetochore tension (40–45). Classic experiments by Bruce Nicklas (46) demonstrated that manually applying tension via a glass microneedle decreases the KMT detachment rate. The Aurora B kinase, a mitotic kinase that localizes to the inner centromere during metaphase in a Haspin kinase dependent manner, is thought to be responsible for mediating tension-dependent stabilization of KMTs (43–45, 47–51). Aurora B regulates many targets including both NDC80c (15, 26, 27, 36, 50, 52–54) and MCAK (55–59). It is not well understood how Aurora B phosphorylation of these targets regulates KMT attachments (34, 38, 47, 51, 60).

Here, we investigate the mechanism of how KMTs detach from the kinetochore and how erroneous KMTs are selectively destabilized. To do so, we first employed a FLIM-FRET assay to characterize NDC80c binding to kinetochores in live cells by measuring the fraction of NDC80c bound to KMTs in cells with a series of NDC80c phosphomutants to perturb NDC80c-KMT binding at kinetochores. We found that the NDC80c binds non-cooperatively to KMTs and that each NDC80c residue that is phosphorylated reduces the NDC80c-KMT binding energy by 0.35 ± 0.03k_B_T. We then employed a photoconversion experiment on these NDC80c phosphomutant cell lines to investigate how perturbing NDC80c binding alters KMT stability. Based on these results, we propose a dual detachment mechanism in which NDC80c binds noncooperatively to KMTs and KMTs detach from the kinetochore when either 1) all NDC80c associated with the KMT spontaneously unbind or 2) following a KMT catastrophe. We also obtained an approximated analytic expression for the relationship between KMT lifetime and fraction of NDC80c bound to KMTs. This dual detachment model is consistent with experiments in cells with phosphomutant NDC80c as well as perturbed MCAK activity. We then investigated how KMTs are destabilized by Aurora B at low-tension, erroneously attached kinetochores, and found that KMT release at low tension is mediated by Haspin-recruited, centromere-localized Aurora B phosphorylation of the NDC80c, which reduces NDC80c affinity for KMTs rather than increasing in KMT catastrophes.

## RESULTS

### The energetics of NDC80c-KMT binding and its modulation by phospho-regulation

To investigate the mechanism by which KMTs detach from the kinetochore, we first sought to better understand the molecular interaction between KMTs and the kinetochore. It is well established that the NDC80c, an outer kinetochore subunit, is the primary coupler between kinetochore microtubules and chromosomes, with roughly 20 NDC80c interacting with each KMT (Figure 1A) (9, 26, 27, 29, 52, 61). The binding affinity of the NDC80c for microtubules is regulated via phosphorylation of 9 sites on the N-terminal tail of the Hec1 subunit of the NDC80c (26, 27, 36, 54, 61, 62). A key question regarding the mechanism of KMT detachment from the kinetochore concerns the energetics of NDC80c-KMT binding and the extent to which NDC80c-NDC80c interactions are cooperative, with prior *in-vitro* studies with purified microtubules and NDC80 subunits giving conflicting results (54, 63). We thus sought to measure the energetics of NDC80c-KMT binding and determine the extent of NDC80c cooperativity at kinetochores *in situ*.

**Figure 1:**
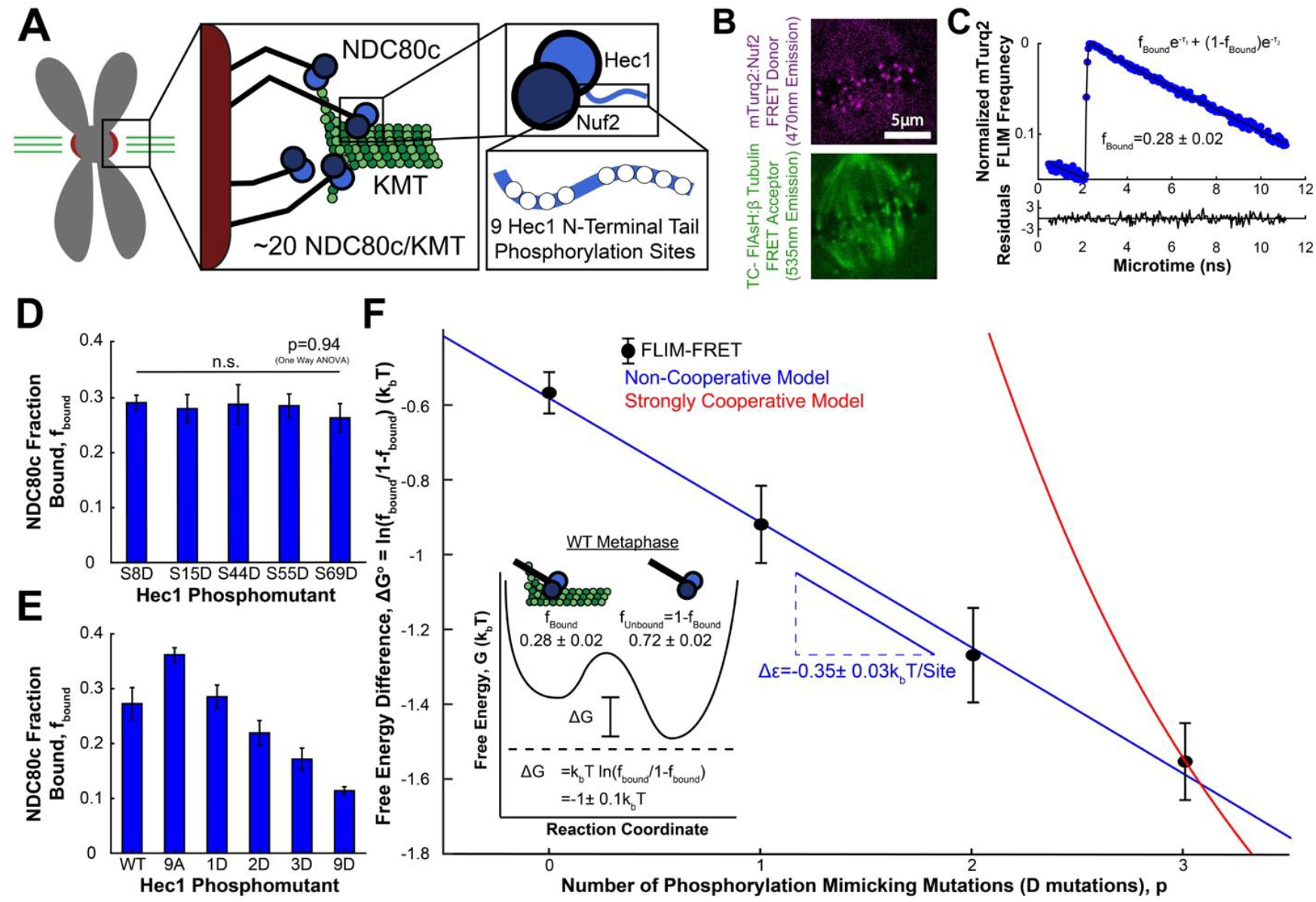
NDC80c binding to KMTs is not cooperative. A) ∼20 NDC80c bind to each KMT at kinetochores. The microtubule binding face of the NDC80c is composed of Nuf2 and Hec1. NDC80c-KMT binding is regulated via phosphorylation of nine phosphorylation sites on the N-terminal tail of Hec1. B) mTurqouise2:Nuf2 FRET donor and the β-tubulin:TC-FlAsH FRET acceptor C) Normalized FLIM histogram of mTurquoise:Nuf2 localized at kinetochores fit a dual-exponential corresponding to the bound and unbound mTurquoise:Nuf2. D) Fraction of NDC80c bound to KMTs with phosphomutant S8D (n=12), S15D (n=12), S44D (n=12), S55D (n=18) and S69D (n=12) Hec1 (error bars are standard error of the mean). E) Fraction of NDC80c bound to KMTs with phosphomutant WT (n=19), 9D (n=17), 3D (n=10), 2D (n=24), 1D (n=18) and 9A (n=24) Hec1. F) Free energy difference between bound and unbound NDC80c at kinetochores vs. the number of phosphorylation mimicking mutations. Black dots: FLIM-FRET experiment. Blue Line: non-cooperative model Red line: strongly cooperative model with interaction energy between NDC80cs set to 0.1k_B_T Inset: Reaction coordiante diagram of NDC80c-KMT binding in metaphase cells. (*f*_*bound*_ =0.28±0.02; Δ*G*^0^=1.0 ± 0.1k_B_T).

We used fluorescence lifetime imaging microcopy (FLIM) – Fluorescence resonance energy transfer (FRET) (64, 65) to measure the fraction of NDC80cs bound to KMTs at kinetochores in metaphase. We employed a previously generated stable U2OS cell line in which the N-terminus of the Nuf2 subunit was fused to an mTurquoise2 FRET donor and the C-terminus of β-tubulin was fused to a TC-FlAsH tag which acts as a FRET acceptor (47). We imaged the cells with two-photon FLIM (Figure 1B) and binned the photons from kinetochores in the mTurquoise2 channel to measure the fluorescence decay of mTurquoise2 at kinetochores (Figure 1C). We found that the fluorescence decay curve of mTurquoise2:Nuf2 at kinetochores showed substantial deviation from a single exponential (Supp. Figure 1) and was well fit to a sum of two exponentials (Figure 1C). This implies that there are two subpopulations of mTurquoise2, which we have previously validated correspond to bound and unbound NDC80c at kinetochores, after correcting the bound fraction to account for only labeling one isotype of β-tubulin (47). Using this procedure on metaphase spindles revealed a mean fraction of *f*_*bound*_= 0.28 ± 0.02 bound and *f*_*unbound*_= 1 − *f*_*bound*_= 0.72 ± 0.02 unbound NDC80cs to KMTs.

We next investigated how phosphorylation of the N-terminal tail of the Hec1 subunit of the NDC80c impacts the association between NDC80 and KMTs in metaphase cells by utilizing a series of phosphomemitic mutants. We created Hec1 mutants in which each of the nine sites was changed to either an alanine (A), to mimic a de-phosphorylated site, or an aspartic acid (D), to mimic a phosphorylated site (26, 27, 37, 53, 54, 61). We inserted an RNAi-resistant, doxycycline inducible transgene of these mutants into the genome, depleted endogenous Hec1 by RNAi, and replaced with mutant Hec1 by addition of doxycycline (35, 47, 53). We first performed kinetochore FLIM-FRET measurements on cells with five different 1D-Hec1 mutants (which all contain one D residue and eight A residues) and found that they exhibit the same fraction of bound NDC80c (p=0.94 ANOVA), indicating that the position of phosphorylated residue does not significantly impact NDC80c-KMT binding (Figure 1D). We then compared mutants mimicking the completely dephosphorylated state (9A-Hec1), and states with one (1D-Hec1), two (2D-Hec1), or three (3D-Hec1) phosphorylated sites, and observed a monotonic decrease in NDC80c fraction bound with increasing number of phosphomemitic mutations (p=2.2×10^−14^ ANOVA) (Figure 1E).

We next to sought to understand the biophysical basis of the observed decrease in NDC80c binding with increasing number of phosphomemitic mutations. In a simple, non-cooperative two-state system in equilibrium, the free energy difference between the two states can be related to the relative mean fraction of the two states via (66):

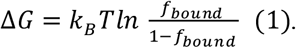

We used Eq. 1 to calculate an apparent free energy difference between bound and unbound NDC80c for each of the phosphomimetic mutants (assuming a simple two-state system) (Figure 1F), which corresponds to the NDC80c-KMT binding affinity, *ε*: i.e. Δ*G* = *ε*. We found that the calculated free energy difference between the bound and unbound states varies linearly with the number of phosphomimetic mutants, suggesting that NDC80c binding is non-cooperative and each additional phosphomimetic site decreasing the free energy of NDC80c binding to KMTs by the same amount: i.e. *ε* = *ε*_0_ + *p*Δ*ε* where *p* is the number of phosphorylated sites, Δ*ε* is the change in the binding affinity from each phosphorylation and *ε*_0_ is the binding affinity of unphosphorylated NDC80c. We next considered an alternative model in which NDC80cs interact with each other to bind cooperatively to KMTs (see Methods). Even if the binding energies between NDC80cs were just one tenth that of the binding energy between NDC80s and KMTs, the predicted trend would drastically depart from the data (Figure 1F, red line; p=0, Bayesian likelihood ratio). A best fit to the full cooperative model gives an interaction energy between NDC80s of −0.006 ± 0.004k_B_T, which is statistically indistinguishable from a model with no cooperativity (Figure 1F blue line; p=0.07, Bayesian likelihood ratio). The fit to the full model also gives Δ*ε*=-0.35 ± 0.03k_B_T/site and *ε*_0_=-0.50 ± 0.02k_B_T, which is statistically indistinguishable from the change in the NDC80c-KMT binding affinity from each phosphorylated residue measured in *in-vitro* single molecule experiments (54) (Δ*ε*=-0.30 ± 0.02k_B_T/site; p=0.11, Bayesian likelihood ratio). Using this model to calculate the free energy difference between bound and unbound NDC80c in wildtype metaphase spindles gives Δ*G*_*WT*_=-1.0 ± 0.1k_B_T (Figure 1F, inset), suggesting that 1-2 residues of the N-terminal tail of the Hec1 are phosphorated in metaphase, which is in agreement with prior measurements of its phosphorylation state (67). Taken together, these FLIM-FRET results indicate that NDC80c at kinetochores bind noncooperatively to KMTs with a binding energy that varies linearly with the number of phosphorylated sites.

### KMTs detach when all the NDC80c spontaneously unbind or following KMT catastrophe

Having characterized the energetics of NDC80c-KMT binding and the (lack of) cooperativity at metaphase kinetochores, we next sought to relate these molecular level biophysical properties to the mechanism of KMT detachment from the kinetochore. To do so, we first measured how the lifetime of KMTs varied with the binding affinity between KMTs and the NDC80c. We utilized a photoconversion assay (12, 68–70) to measure the lifetime of KMTs in phosphomutant Hec1 cells. We generated a U2OS cell line with mEOS3.2:α-tubulin to mark microtubules and SNAP-SiR:hCentrin2 to mark the spindle poles. mEOS3.2 is a photoconvertible fluorescent protein, which switches from green to red emission after stimulation with 405nm UV light. We used a home-built photoconversion system with a 405nm diode laser to photoconvert a line of tubulin across the spindle perpendicular to the spindle axis (Figure 2A) (12, 71). We then projected the photoconverted intensity onto the spindle axis (Figure 2B) and fit the profile to a Gaussian to measure the height of the peak (Figure 2C). We fit the height of the photoconverted peak, corrected for background tubulin incorporation and bleaching (Supp. Figure 2), to a dual-exponential decay (Figure 2D). The slow-decay lifetime of this dual exponential decay is typically attributed to the slow turnover of the stabilized KMTs, consistent with recent results from mitotic human tissue culture cells showing that the fraction of tubulin in KMTs in electron tomography reconstructions matches the slow-decay fraction of the decay after photoconversion (12). Using this photoconversion assay, we measured the KMT lifetime in Hec1 phosphomutant cells (Supp. Figure 3) and found that KMT stability increased with the number of de-phosphomimetic mutations, with KMTs more stable in 9A-Hec1 cells than 9D-Hec1 cells (Figure 2E. 9D: 2.6±0.4min, 9A: 4.0±0.4min, p =0.014, Student’s t-test).

**Figure 2:**
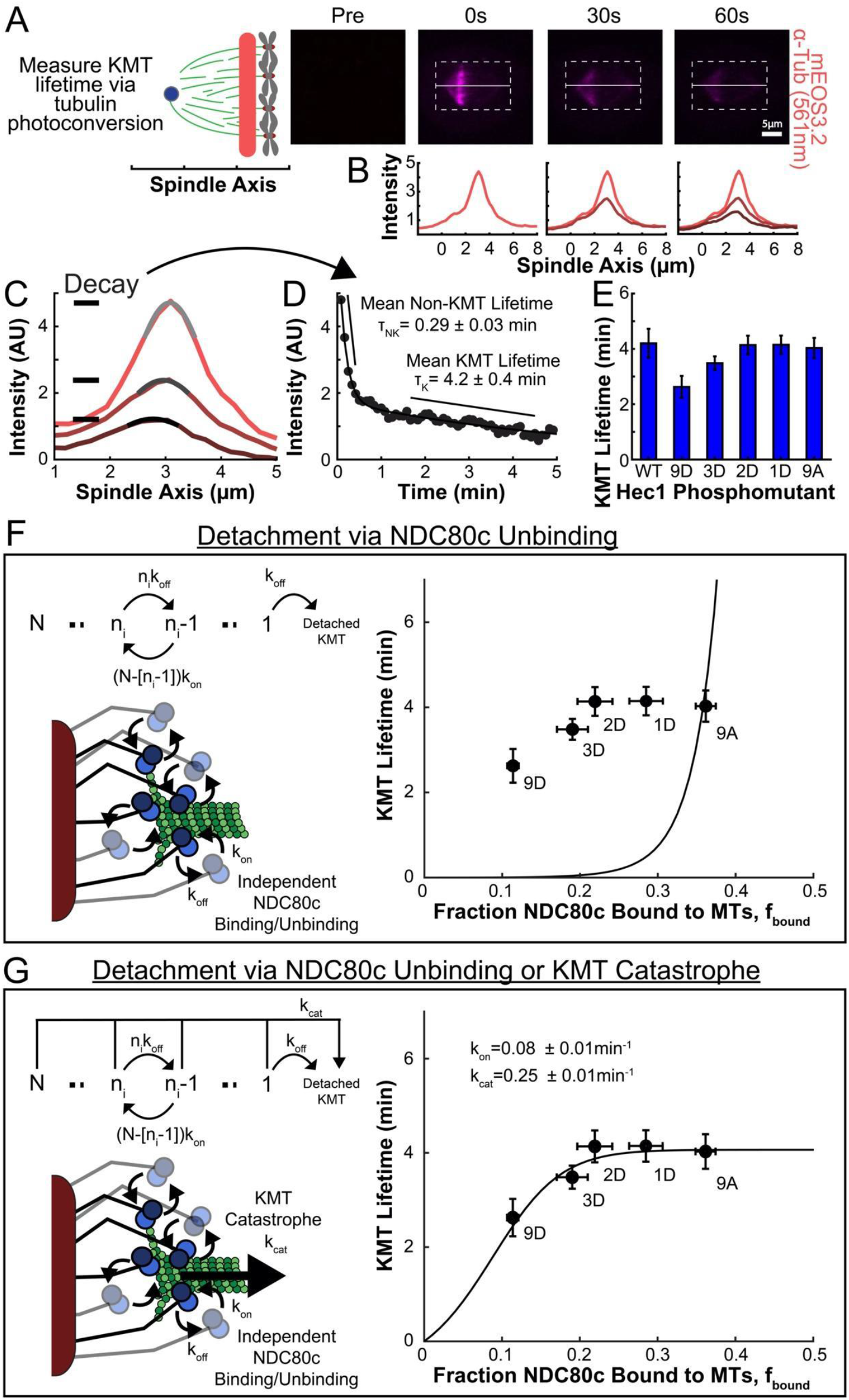
Mechanism of KMT detachment. A) Images from a photoconversion experiment showing post-conversion mEOS3.2:α-tubulin (561nm, magenta). Images are shown immediately preceding, 0, 30, and 60s after photoconversion with a focused 405nm diode laser B) Line profile generated by averaging 3.5μm on either side of the spindle axis in dotted box in A. C) Line profiles (shades of red) fit to Gaussian profiles (shades of gray). Lighter shades are earlier times. (D) Black dots: Fit intensity of the line profile over time, corrected for background and bleaching. Black line: dual-exponential fit decay. E) Bar plot of mean KMT lifetimes for cells expressing phosphomutant Hec1 F) Left: A rate transition model with states corresponding to the number of bound NDC80c with an additional state with the KMT detached and NDC80c bind and unbind independently. Transitions between states are defined only by NDC80c binding: *n*_*i*_ → *n*_*i*−1_: *n*_*i*_*k*_*off*_; *n*_*i*−1_ → *n*_*i*_: *n*_*i*−1_ *k*_*on*_. The KMT detaches when the last NDC80c unbinds at rate *k*_*off*_. Right: KMT Lifetime vs Fraction of NDC80c bound to MTs for Hec1 phosphomutants (black dots) compared with model predictions (black line). G) Left: A model with independent NDC80c binding and unbinding and KMT catastrophes. Transitions between states defined by NDC80c binding: *n*_*i*_ → *n*_*i*− 1_: *n*_*i*_*k*_*off*_; *n*_*i*−1_ → *n*_*i*_ : *n*_*i*−1_*k*_*on*_ and by KMT catastrophe: *n*_*i*_ → *Detached*: *k*_*cat*_. Right: KMT Lifetime vs Fraction of NDC80c bound to MTs for Hec1 phosphomutants (black dots) compared with model predictions (black line)

**Figure 3:**
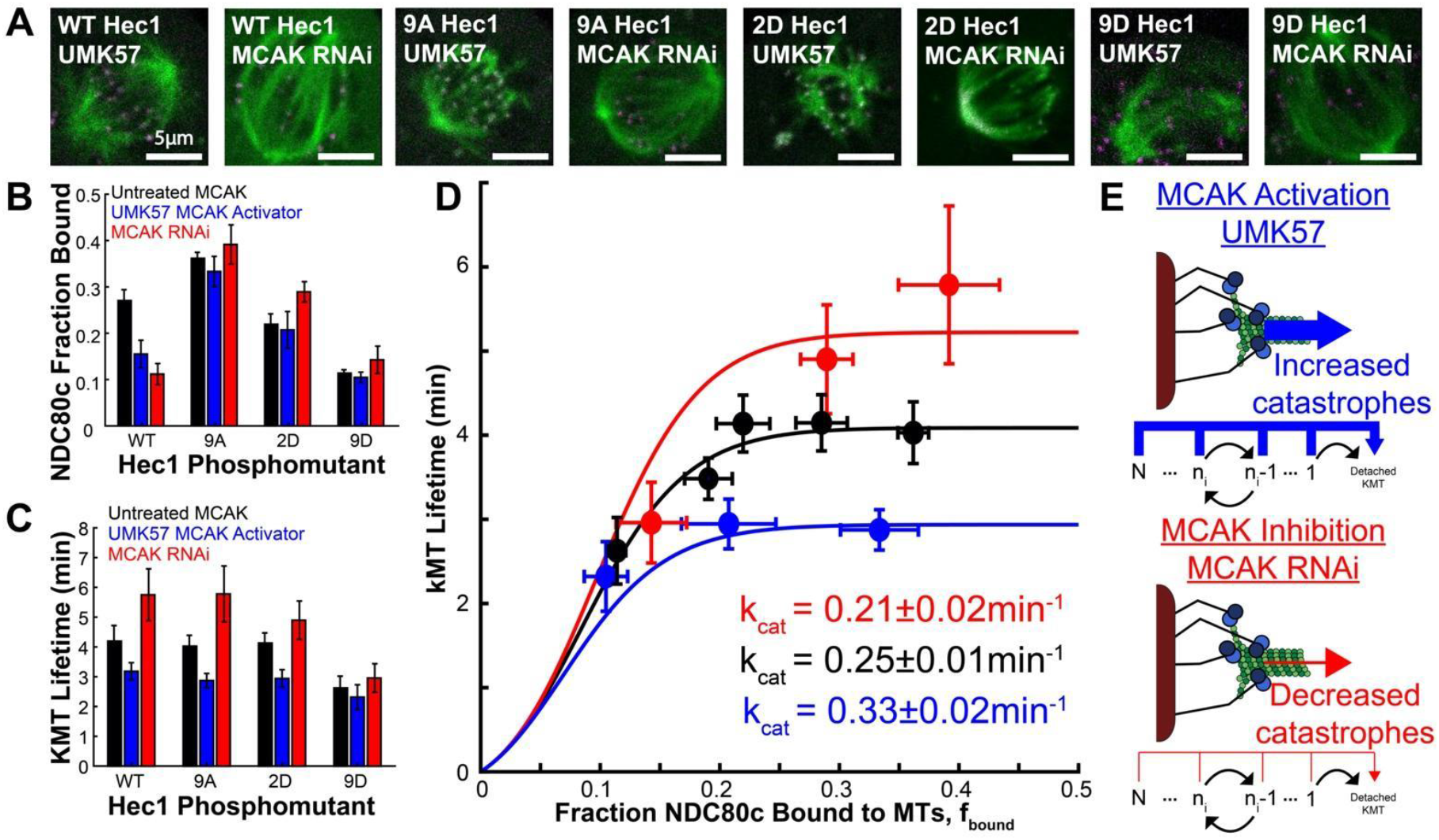
Both MCAK hyperactivation and depletion alter the KMT catastrophe rate. A) UMK57 treated and MCAK depleted cells with WT-,9A-,2D- and 9D-Hec1. B) NDC80c fraction bound for wild type MCAK (black), UMK57 treated hyperactivated MCAK (blue) and MCAK depleted (red) cells with WT-,9A-,2D- and 9D-Hec1 (error bars are standard error of the mean. Untreated: WT-Hec1 n=19, 9D-Hec1 n=17, 2D-Hec1 n=24, 9A-Hec1 n=24; MCAK RNAi: WT-Hec1 n=7, 9D-Hec1 n=7, 2D-Hec1 n=11, 9A-Hec1 n=11; UMK57 MCAK: WT-Hec1 n=12, 9D-Hec1 n=7, 2D-Hec1 n=12, 9A-Hec1 n=10). C) KMT lifetime for wild type MCAK (black), UMK57 treated hyperactivated MCAK (blue) and MCAK depleted (red) cells with WT-,9A-,2D- and 9D-Hec1. (Untreated: WT-Hec1 n=22, 9D-Hec1 n=41, 2D-Hec1 n=32, 9A-Hec1 n=40; MCAK RNAi: WT-Hec1 n=16, 9D-Hec1 n=17, 2D-Hec1 n=21, 9A-Hec1 n=20; UMK57 MCAK: WT-Hec1 n=33, 9D-Hec1 n=19, 2D-Hec1 n=27, 9A-Hec1 n=31) D) KMT lifetime vs fraction of NDC80c bound to KMTs for cells expressing Hec1 phosphomutants with wild type (black dots) hyperactivated (blue dots) and depleted (red dots) MCAK fit to the dual detachment mechanism model (red, blue and black lines). E) MCAK hyperactivation with UMK57 increases the KMT catastrophe rate while MCAK inhibition with MCAK RNAi decreases the KMT catastrophe rate.

We next developed a biophysical model of how KMTs detach from the kinetochore to connect the KMT lifetimes observed in the phosphomutant photoconversion experiments with the NDC80c bound fractions observed in the phosphomutant FLIM-FRET experiments. We considered models in which there are a fixed number, *N*, of NDC80s per KMT, each of which is either bound or unbound to the KMT, and in which the KMT detaches from the kinetochore when no NDC80cs are bound to it (Figure 2F, left). Our Hec1 phosphomutant photoconversion and FLIM-FRET experiments indicated that perturbing NDC80c binding kinetics altered the KMT lifetime, so we initially explored a model where only NDC80c binding kinetics regulated the KMT lifetime. Motivated by our FLIM-FRET results that NDC80c binding to KMTs was not cooperative, we assumed that *N* NDC80cs independently bind and unbind to KMTs and defined *N* + 1 total states with *n* = 0, 1,…, *N* NDC80c bound (Figure 2F, top left panel). The transition rate from the *n*^th^ state with *n* NDC80c bound to the (*n* − 1)^th^ state with *n* − 1 NDC80c bound is given by the rate that any of *n* bound NDC80c unbind from the KMT, *nk*_*off*_, and the transition rate from the (*n* − 1)^th^ state to the *n*_*i*_ state is given by the rate that any of the (*N* − (*n* − 1)) unbound NDC80c bind to the KMT, (*N* − (*n* − 1))*k*_*on*_. In this model, a KMT detaches from the kinetochore after that last NDC80s unbinds it and the system enters the *n* = 0 “detached” state.

To determine whether this model, in which KMT detachment is determined solely by NDC80c binding kinetics, was consistent with both the NDC80c bound fraction and the KMT lifetime experiments, we calculated the distribution of KMT lifetimes as a function of NDC80c bound fraction (see *Supplemental Methods, Section 1*). In this model, the KMT lifetime corresponds to the mean first passage time to the “detached KMT” state. For relevant parameters, the first passage time to the *n* = 0 “detached KMT” state is dominated by the last step (i.e. the unbinding of the last NDC80c transition from the *n* = 1 to the *n* = 0 state), in which case the KMT lifetimes can be analytically calculated to be approximately exponential distributed with a mean of:

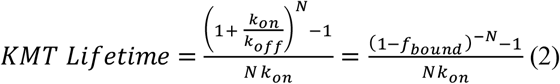

Where the equality results from using the relationship 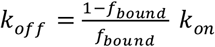. Based on prior measurements of the number of NDC80c (∼200 NDC80c/kinetochore (29)) and the number of KMTs (∼10 KMTs/kinetochore (9)) at each kinetochore, we set *N* = 20 NDC80c per KMT. Thus, there is only a single unknown parameter in the model to be fit to the phosphomutant data, *k*_*on*_. A best fit to this model predicts a sharp increase in KMT lifetime with increasing *f*_*bound*_, in clear disagreement with experimental measurements (Figure 2F, right). This discrepancy between the model and measurements is not a consequence of assuming noncooperative NDC80c binding: A model where all NDC80c bind cooperatively as a single binding unit (which is formally equivalent to the previous model but with *N* = 1) predicts a similar increase in the KMT lifetime with increasing fraction bound, leading to a similarly poor fit (Supp. Figure 4). Thus, a model in which KMTs detach only when all NDC80c unbind is not consistent with the observed KMT lifetime and NDC80c fraction bound in the Hec1 phosphomutants.

**Figure 4:**
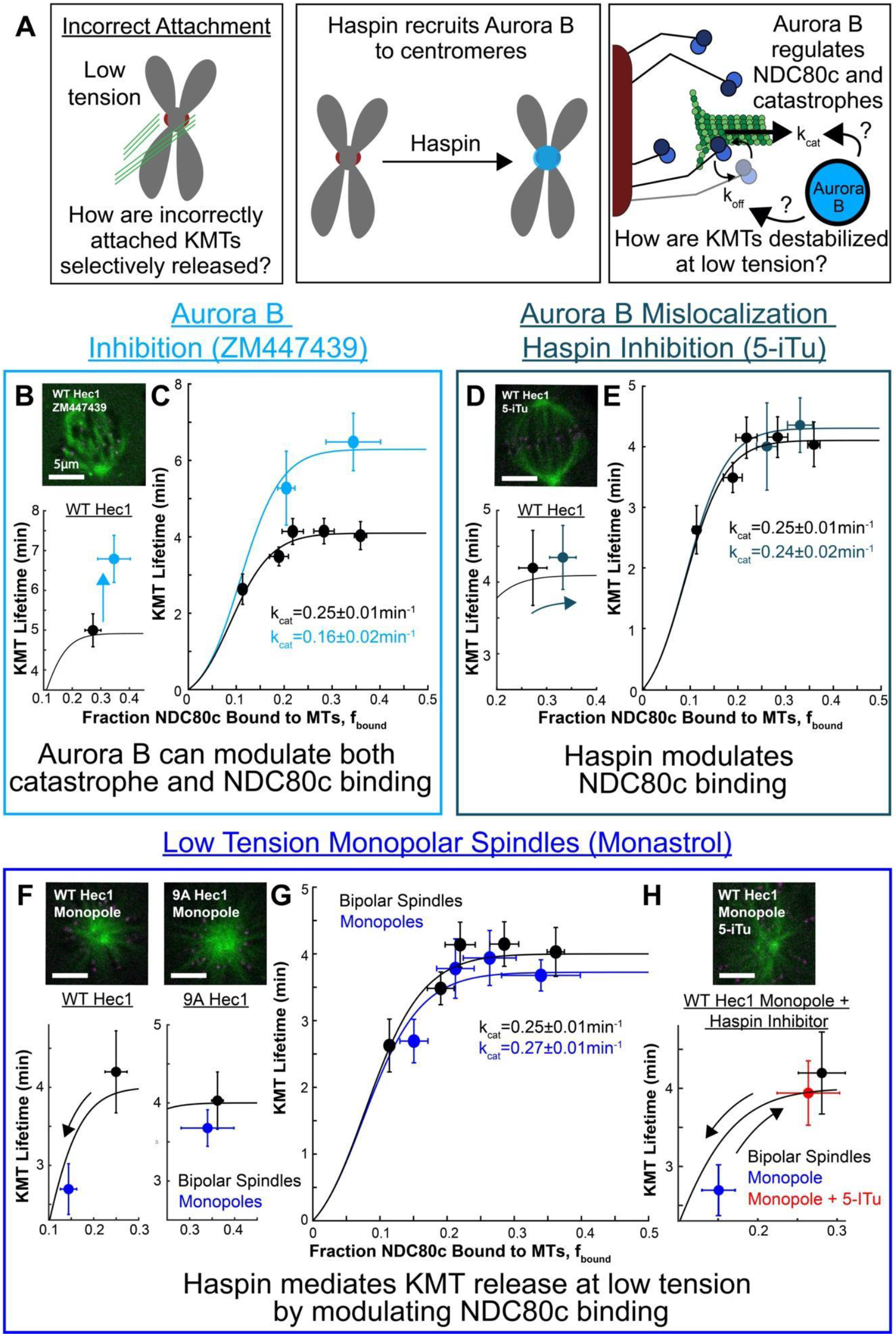
Low-tension KMTs are destabilized by a Haspin-mediated increase in the NDC80c unbinding k_off_ rate. A) Left: Schematic depicting a chromosome with low inter-kinetochore tension with both kinetochores attached to the same spindle pole. Middle: Haspin recruits the Aurora B kinase to the inner centromere by phosphorylating the H3T3 histone. Right: Aurora B can regulate either the NDC80c unbinding rate or the KMT catastrophe rate B) Sample image, KMT lifetime and NDC80c bound fraction of cells expressing WT-Hec1 and treated with ZM447439 Aurora B inhibitor (cyan) compared to untreated wild type (black) C) KMT lifetime and NDC80c bound fraction of WT- and 2D-Hec1 phosphomutants fit to the dual detachment mechanism for ZM447439 Aurora B inhibited cells (cyan, *k*_*cat*_=0.16±0.02min^-1^). D) Sample image, KMT lifetime and NDC80c bound fraction of cells expressing WT-Hec1 and treated with 5-ITu Haspin inhibitor compared to wild type E) KMT lifetime and NDC80c bound fraction of WT- and 2D-Hec1 phosphomutants fit to the dual detachment mechanism for 5-ITu Haspin inhibited cells (*k*_*cat*_=0.14±0.02min^-1^). F) Sample image, KMT lifetime and NDC80c bound fraction of monopolar spindles (treated with 200µM monastrol) expressing WT- and 9A-Hec1 compared to wild type bipolar spindles G) KMT lifetime and NDC80c bound fraction of monopolar phosphomutants fit to the dual detachment mechanism (*k*_*cat*_=0.27±0.02min^-1^). H) Sample image, KMT lifetime and NDC80c bound fraction of monopolar spindles expressing WT-Hec1 treated with 5-ITu Haspin inhibitor compared to wild type monopolar and bipolar spindles.

Prior work has demonstrated that both NDC80c phosphorylation (52–54, 65) and the activity of microtubule catastrophe factors (25, 32, 33, 56, 57, 72) can regulate the lifetime of KMTs. We therefore sought to incorporate the impact of catastrophe factors into a model of KMT detachment. We proposed that, in addition to KMTs detaching due to NDC80c unbinding, KMTs can also catastrophically detach at a constant rate *k*_*cat*_ that is independent of NDC80 binding (Figure 2G, left). In this model, the KMT can detach when either all NDC80c spontaneously unbind from the KMT or after the KMT undergoes a catastrophe. We then analytically calculated the KMT lifetimes predicted by this model (see *Supplemental Methods, Section 3*) and found that, in the relevant parameter regime, they are exponentially distributed (consistent with the photoactivation experiments), with a mean lifetime approximately given by:

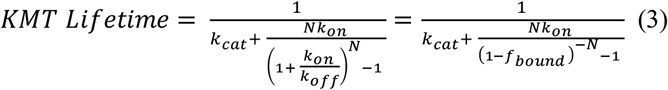

Where we again used the relationship 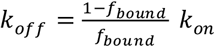. This model with both NDC80c unbinding and KMT catastrophes fit well to the Hec1 phosphomutant data (*k*_*on*_ =0.08±0.01min^-1^ and *k*_*cat*_=0.25±0.01min^-1^) (Figure 2G, right). We confirmed that the analytic solution (equation 3) was a good approximation at the fit model parameters by computing the distribution of KMT lifetimes, fraction of NDC80c bound and kinetochore occupancy numerically (see *Supplemental Methods, Section 4*). We conclude that a dual-detachment model, where KMTs detach when either all the NDC80c unbind from the KMT or following catastrophic detachment, is consistent with both the KMT lifetime and the NDC80c binding observed in the Hec1 phosphomutants.

To further test this proposed dual-detachment model, we investigated the impact of perturbing MCAK, a known catastrophe factor (25, 30–33, 59). We used the knockdown-rescue protocol to investigate cells with WT Hec1 and 9A, 2D, and 9D Hec1 phosphomutants that were either subject to MCAK RNAi knockdown or treated with UMK57, a small molecule MCAK potentiator (32, 33, 73, 74). (Figure 3A). We performed FLIM-FRET under these different conditions to measure the impact on NDC80c binding (Figure 3B) and photoconversion to measure KMT lifetimes (Figure 3C). We combined these two datasets by plotting KMT lifetime vs NDC80c bound fraction and fit the resulting curve with the predictions from the dual-detachment model (Figure 3D). Both the MCAK RNAi and UMK57 data fit well to the dual detachment model, with the same NDC80c binding rate as control cells (*k*_*on*_ =0.08±0.01min^-1^), but a lower KMT catastrophe rate than wildtype in the MCAK RNAi cells and a higher KMT catastrophe rate than wildtype in the UMK57 cells (siMCAK: *k*_*cat*_=0.21±0.02min^-1^, UMK57: *k*_*cat*_=0.33±0.02min^-1^, p=2×10^−6^ Bayesian parameter comparison). Thus, hyperactivating MCAK with UMK57 increases the catastrophe rate and inhibiting MCAK activity by RNAi knockdown decreases the catastrophe rate, both without modifying NDC80c-KMT interactions (Figure 3E). This behavior is consistent with prior work demonstrating that MCAK is a catastrophe factor that can modulate KMT stability (25, 30–33, 59, 73, 74), and further supports the validity of the dual-detachment model in which KMTs detach from the kinetochore due to either: 1) all NDC80cs spontaneously unbinding or 2) catastrophic detachment. Furthermore, this model provides a means to quantitively relate changes in the NDC80c-KMT binding energy to changes in KMT stability: Using Equation 3 with the relationship 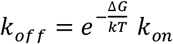, and the result that Δ*G* = *ε* = *ε*_0_ + *p*Δ*ε*, gives a quantitative prediction for how KMT lifetime depends on the catastrophe rate, *k*_*cat*_, the number of NDC80c molecules per kinetochore microtubule, *N*, and the number of phosphorylated residues on NDC80c, *p*:

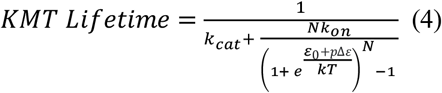

### Haspin-localized Aurora B destabilizes low-tension KMTs by weakening NDC80c binding not regulating KMT catastrophes

Having developed a dual-detachment model which relates both the energetics of NDC80c-KMT binding and KMT catastrophe to KMT detachment, we wanted to investigate the mechanism by which erroneous KMTs are selectively removed from kinetochores during mitotic error correction. Classic experiments by Bruce Nicklas demonstrated that manually applying tension across non-bioriented kinetochores stabilizes the attached KMTs (46). This tension-dependent stabilization is thought to mediate the selective removal of KMTs from erroneously attached kinetochores, with KMTs destabilized at low-tension, unaligned, incorrectly attached kinetochores (40–45). Subsequent work has suggested that this process is mediated by the Aurora B kinase (48–51, 60, 75), which localizes to inner centromeres during metaphase in a Haspin kinase dependent manner (76–78). Aurora B phosphorylates many targets, including both MCAK (55–59, 73) and the N-terminal tail of Hec1 in the NDC80c (26, 27, 44, 47, 48, 54) meaning that it is possible that Aurora B could regulate either the NDC80c unbinding detachment mechanism or the KMT catastrophe detachment mechanism.

We first inhibited Aurora B with the drug ZM447439 (79) and performed both photoconversion experiments and FLIM-FRET experiments. The KMT lifetime significantly increased after Aurora B inhibition (p=0.03, Students’ t-test) resulting in the Aurora B inhibited point falling off the dual-detachment model curve of WT cells and indicating that inhibiting Aurora B altered the KMT catastrophe rate (Figure 4B). The increase in the NDC80c fraction bound after Aurora B inhibition was not statistically significant (p=0.3, Students’ t-test), which is surprising because Aurora B is known to phosphorylate NDC80c (26, 27, 44, 47, 48, 54). We speculate the explanation for the lack of impact of Aurora B inhibition on NDC80c binding in metaphase is because, at this time in mitosis, the wild type NDC80c bound fraction is similar to the almost fully de-phosphorylated 1D-Hec1 mutant (p=0.97; ANOVA), so Aurora B inhibition likely does little to alter the bound fraction of the already mostly de-phosphorylated wild type metaphase NDC80c. Consistent with this hypothesis, previous work showed that inhibiting Aurora-B in prometaphase, when NDC80c binding is lower than in metaphase, does lead to a significant increase in NDC80c binding (47). We next performed photoconversion and FLIM-FRET measurements on ZM447439 treated, Hec1 phosphomutant cells and fit the dual detachment model, giving *k*_*cat*_ = 0.16±0.02min^-1^ (Figure 4C), which is significantly less than the catastrophe rate in WT cells (p=1×10^−4^ Bayesian likelihood ratio). Thus, global inhibition of Aurora B in metaphase primarily stabilizes KMT detachments by reducing catastrophes. To explore the role of the centromere localized Aurora B pool, thought to be responsible for conferring tension-dependent KMT stabilization, we next disrupted Aurora B localization to centromeres with 5-Itu (76) and performed photoconversion experiments and FLIM-FRET experiments. The 5-ITu data remained on the dual-detachment model curve of WT cells (Figure 4D). We took photoconversion and FLIM-FRET measurements of 5-ITu treated, Hec1 phosphomutant cells and fit the dual detachment model, giving *k*_*cat*_=0.24±0.02min^-1^, which is indistinguishable from the catastrophe rate in WT cells (p=0.82, Bayesian likelihood ratio) (Figure 4E). Taken together, this indicates that Aurora B regulates both KMT catastrophes and NDC80c binding to KMTs, but the pool of Aurora B recruited to centromeres by Haspin only regulates NDC80c binding.

To investigate this further, we performed experiments in monopolar spindles, generated by inhibiting kinesin-5 with a small molecule inhibitor, Monastrol, where kinetochores are not bioriented and are under low tension (47, 80, 81). We measured the KMT lifetime and fraction of NDC80c bound to KMTs in the low tension, monopolar spindles and found that both the KMT lifetime (p=8×10^−4^ Students’ t-test) and fraction of NDC80c bound (p=2×10^−3^ Students’ t-test) were lower in monopolar spindles than WT bipolar spindles, but the monopolar spindle datapoint still lay on the same fit dual-detachment model curve as WT bipolar spindles (Figure 4F). This suggests that the low tension KMTs are destabilized by weakened NDC80c binding to KMTs. We measured the KMT lifetime and NDC80c bound fraction in monopolar spindles expressing a 9A-Hec1 phosphomutant and found that they had the same NDC80c bound fraction and KMT lifetime as biopolar 9A-Hec1 spindles (NDC80 bound fraction: p=0.60, Students’ t-test; KMT lifetime: p=0.61, Students’ t-test), arguing that the tension dependency of KMT detachments requires the phosphorylation of the Hec1 N-terminal tail. Combining the data from monopolar spindles with WT Hec1 and monopolar spindle with Hec1 phosphomutants and fitting to the dual-detachment model gave a KMT catastrophe rate of *k*_*cat*_ = 0.27±0.01min^-1^ in monopolar spindles, statistically indistinguishable from the KMT catastrophe rate in bipolar spindles (p=0.24, Bayesian likelihood ratio) (Figure 4G). We next investigated the contribution of centromere localized Aurora B to destabilizing low tension KMT detachments by treating monopolar spindles with the 5-ITu Haspin inhibitor. We performed photoconversion experiments and FLIM-FRET experiments, which revealed that the addition of 5-ITu to monopoles caused an increase in both NDC80c fraction bound and KMT lifetimes relative to monopoles without 5-ITu (NDC80 bound fraction: p=0.02, Students’ t-test; KMT lifetime: p=0.02, Students’ t-test), to values which were indistinguishable to those of WT bipolar spindles (NDC80 bound fraction: p=0.31 Students’ t-test; KMT lifetime: p=0.65, Students’ t-test) (Figure 4H). This indicates that centromere-localized Aurora B is required for destabilizing KMTs at low tension. Taken together, these results indicate that low tension KMT detachments are destabilized due to phosphorylation of the Hec1 tail by Haspin-localized Aurora B kinase reducing the affinity of NDC80c to KMTs (without modulation of KMT catastrophes).

## DISCUSSION

The removal of incorrectly attached KMTs is crucial for the correction of mitotic errors and proper chromosome segregation; however, the biophysical basis of ND80c binding to KMTs to the kinetochore, the mechanism by which KMTs detach from the kinetochore, and how KMTs are selectively destabilized at erroneously attached chromosomes, is not well understood. Our results indicate that individual NDC80c bind noncooperatively to KMTs and support a dual-detachment mechanism where individual NDC80c independently bind and unbind to KMTs and KMTs detach either when all the associated NDC80c spontaneously unbind or following a KMT catastrophe. In this model, KMT stability can be regulated by altering either the rate that NDC80c unbind from KMTs or the KMT catastrophe rate. We found that Aurora B kinase regulated both NDC80c binding and KMT catastrophes, though disrupting Aurora B localization to the inner centromere by inhibiting the Haspin kinase only impacted NDC80c binding. In monopolar spindles where chromosomes are erroneously attached and subject to low tension, we found that NDC80c binding was perturbed but the KMT catastrophe rate was similar to wild - type bipolar metaphase spindles. Disrupting Aurora B localization to inner centromeres by inhibiting Haspin kinase with 5-ITu restored the NDC80c binding and KMT stability of monopolar spindles to the levels they exhibit in wild-type bipolar metaphase spindles. Taken together, these results support a model in which KMTs at erroneously attached chromosomes are destabilized by Haspin-localized Aurora B kinase phosphorylating Hec1 and thereby regulating NDC80c binding to KMTs.

There has been substantial work characterizing NDC80c-KMT binding *in* vitro (41, 45, 54, 63, 82, 83); however, little is known about NDC80c-KMT binding at kinetochores in live cells. We found that the unbound NDC80c state is favored (Δ*G*=-1.0 ± 0.1k_B_T) over the bound NDC80c state in wildtype metaphase cells. The structural basis for this favorability of the unbound state is unclear, as the interactions between the NDC80c and tubulin are almost certainly favorable (82). One possibility is that the linker between the NDC80c and the inner kinetochore is relatively flexible, so the unbound NDC80c is able to diffuse in the vicinity of the KMT, giving a large number of microstates for the unbound NDC80c to sample and entropically favoring the unbound state over the rigid bound state. Alternatively, the linker between the NDC80c and the inner kinetochore could be very rigid and NDC80c binding to KMTs could induce unfavorable conformational changes elsewhere in the kinetochore. Structural studies of the kinetochore in cells would be informative in further understanding NDC80c-KMT binding.

On the key question of whether NDC80 binding is cooperative, *in vitro* studies of NDC80c binding to microtubules in solution have found conflicting results (54, 63). Our results on NDC80c binding to KMTs in spindles indicates that the cooperative interaction energy between bound NDC80c subunit is statically indistinguishable from zero and much weaker than the free energy difference between the bound and unbound wild-type NDC80c, indicating the NDC80c binding to KMTs is not cooperative *in situ*. Additionally, the change in binding affinity from each phosphorylated residue we measured at kinetochores (Δ*ε*/Site=-0.35 ± 0.03k_B_T/Site) was statistically indistinguishable from the change measured in *in-vitro* single molecule experiments (54) (Δ*ε*/Site=-0.30 ± 0.02k_B_T/Site; p=0.11, Bayesian likelihood ratio). In these single molecule experiments, individual NDC80c are visualized binding and unbinding to microtubules, so changes in binding affinity can be measured neglecting cooperative interactions between NDC80cs. The agreement between the Δ*ε* measured at kinetochores and *in-vitro* single-molecule results, where cooperativity can be neglected, further supports that the binding between NDC80c in non-cooperative.

Our results support a model where individual NDC80c independently bind and unbind to KMTs and KMTs detach when either: 1) All NDC80c spontaneously unbind from the KMT or 2) Following a KMT catastrophe (Figure 5A, upper panel). The fit NDC80c unbinding rate in wildtype metaphase spindles was 0.26 ± 0.04 min^-1^, meaning that an NDC80c remains bound to a KMT for 3.9 ± 0.7 min (Figure 5A, lower panel). Our tubulin photoconversion measurements indicate that metaphase KMTs remain attached to the kinetochore for an average of 4.2 ± 0.4 min. Tubulin dimers incorporate at the plus end of KMTs and flux away from the kinetochore along the walls of the KMTs by subsequent tubulin polymerization at the plus end at a rate of ∼1 μm/min (12, 69, 70) so over the ∼4 minutes that an NDC80c remains bound to a KMT, a tubulin dimer will move ∼4 μm from the kinetochore. Since the NDC80c remains localized at the kinetochore, the NDC80c must transition from binding one tubulin dimer to another tubulin dimer to remain bound to the KMT, perhaps diffusing along the surface of the microtubule as proposed in prior work (54, 84, 85).

**Figure 5:**
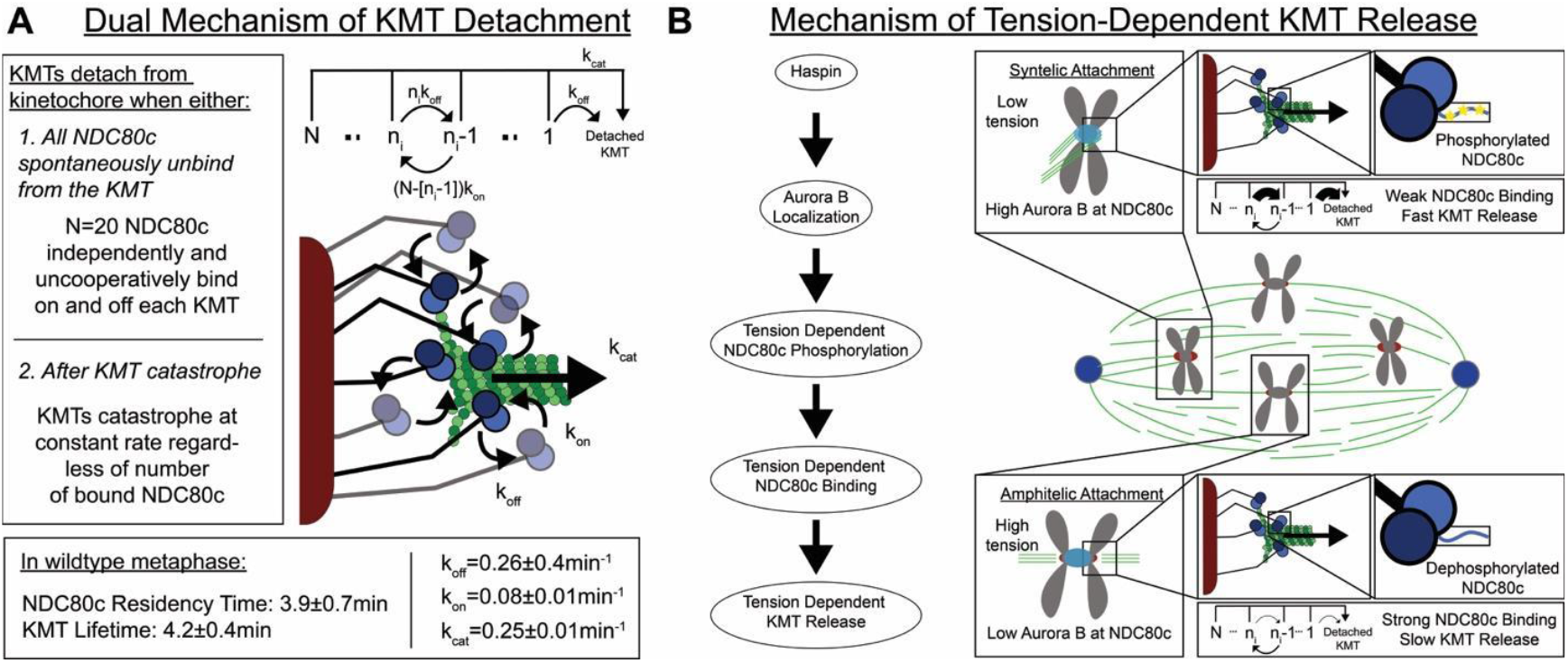
Summary of KMT Detachment Regulation. A) NDC80c at metaphase kinetochores bind independently and uncooperatively to KMTs. KMTs detach from the kinetochore when either: 1. All the NDC80cs spontaneously unbind or 2) Following a KMT catastrophe. The NDC80c bind to MTs at rate *k*_*on*_ ∼0.1min^-1^ and unbind from KMTs at rate *k*_*off*_ ∼0.3min^-1^. The KMT catastrophe rate *k*_*cat*_∼0.25min^-1^ The NDC80c remain bound to the KMT for an average of 4 minutes. KMTs remain attached to kinetochores for an average of ∼4 minutes. B) KMTs are destabilized at low tension by recruitment of Aurora B to centromeres by Haspin. Centromere-localized Aurora B phosphorylates NDC80c in a tension-dependent manner, which confers tension dependence to NDC80c binding and KMT release. Sister kinetochores with incorrect attachments that are not bioriented are under low tension with highly phosphorylated NDC80c, weak NDC80c binding to KMTs and fast KMT release. Sister kinetochores with correct amphitelic attachments are bioriented and under high tension with dephosphorylated NDC80c, strong NDC80c to KMTs and slow KMT release.

We also derived a simplified analytical expression describing the relationship between KMT lifetime and the fraction of NDC80c bound to KMTs in the dual detachment model, using appropriate approximations. The resulting formula indicates that in parameter regimes with a high bound fraction, the KMT lifetime is primarily determined by the catastrophic detachment rate. In contrast, when the bound fraction is low, both detachment mechanisms contribute. Interestingly, this analytical framework, based on first-passage time calculations, has also been applied to chromosome seggregation errors and error correction at a larger scale (4), revealing a self-similar property of the behavior of the spindle and its constituents.

Selective stabilization of KMTs attached to bioriented chromosomes is crucial to ensure accurate chromosome segregation and is thought to be mediated by an increase in inter-kinetochore tension upon biorientation and by the Aurora B kinase, a key mitotic kinase that is localized to the inner centromere by the Haspin kinase. The mechanism of how increased inter-kinetochore tension confers stability to KMTs, is not well understood, though the higher concentration of Aurora B kinase at low-tension kinetochores than at high-tension kinetochores is thought to be crucial for conferring tension-dependent KMT stabilization (26, 27, 40–45, 51, 56, 57, 75, 86). By fitting data from monopolar spindles, where chromosomes cannot biorient and kinetochores are at low tension (47, 70, 77, 78, 86–88), to the dual detachment model, we found that NDC80c binding to KMTs was weaker in monopolar spindles than in bipolar spindles, but the KMT catastrophe rate was similar (Figure 4), indicating that tension dependent KMT stabilization is mediated by regulating NDC80c binding. Inhibiting Aurora B phosphorylation of the NDC80c, either by expressing 9A-Hec1 phosphomutants that Aurora B cannot phosphorylate or by decreasing the concentration of Aurora B at kinetochores by inhibiting the Haspin kinase, prevented KMT lifetime and NDC80c binding from decreasing in monopolar spindles (relative to bipolar spindles), indicating that tension-dependent regulation of NDC80c binding is mediated by Aurora B phosphorylation of NDC80c. Taken together, these observations are consistent with a model for destabilizing erroneous, low tension KMT attachments based on phosphoregulation of NDC80c-KMT binding by Aurora B: the concentration of Aurora B at kinetochores is regulated by tension in a Haspin dependent manner, such that low-tension kinetochores are associated with higher concentrations of Aurora B, which leads to more NDC80c phosphorylation, weaker NDC80c binding to KMTs and destabilized KMTs (Figure 5B).

The experiments and modeling we present here argue that KMTs at erroneously attached kinetochores are destabilized by phosphorylation of nine residues on the N-terminal tail of the Hec1 subunit of the NDC80c by the Aurora B kinase at low-tension kinetochores. Through bottom-up quantification over a range of length scales, we have elucidated how phosphorylation of the Hec1 N-terminal tail alters NDC80c binding to KMTs, how this altered NDC80c-KMT binding destabilizes KMTs and finally how the phosphorylation of these residues is regulated by Aurora B to selectively destabilize KMTs at low-tension, erroneously attached kinetochores. This modeling explains the biophysical basis of mitotic error correction at the level of phosphorylation of specific residues on the Hec1 N-terminal tail, connecting the molecular interaction of the NDC80c and KMTs to accurate chromosome segregation. Further testing this proposal will require careful quantification of the mitotic error rate in response to perturbed KMT lifetime and perturbed Aurora B activity, which will be an exciting challenge for future experiments.

One core aim of cell biology is understanding how the behavior of cells arises from the molecular interactions of proteins. There has been significant interest in developing thermodynamics models of protein interactions in cells to explain, for example, regulation of gene expression (89, 90), dynamics of cytoskeletal filaments and molecular motors (91, 92), kinetic proofreading (93), the formation of condensates (94), and cellular metabolic fluxes (95, 96); however, there has been little work done to validate these models at the molecular level by measuring binding energies and cooperativity *in-situ*. We directly measured the free energy of NDC80c-KMT binding at kinetochores *in-situ* and, based on these measurements, developed and validated a model of how KMTs detach from the kinetochore by measuring KMT detachment rates *in-situ*. Based on this model, and measurements of NDC80c binding and KMT detachment at erroneously attached kinetochores, we demonstrated that erroneous KMTs are selectively destabilized by altering a single thermodynamic parameter: the free energy of binding between the NDC80c and KMTs. *In-situ* quantitative measurements of, for example, transcription factor binding, molecular motors walking along cytoskeletal filaments, cellular metabolic fluxes would be similarly informative for building and validating biophysical models to explain the thermodynamic basis of cellular behavior.

## MATERIALS AND METHODS

### Cell Culture and Cell Line Generation

U2OS cells were acquired from ATCC and cultured in DMEM (ThermoFisher) supplemented with 10% tetracycline-free FBS (ThermoFisher) and Pen-Strep (ThermoFisher) in a humified incubator with 5% CO_2_ maintained at 37°C. A stable cell line expressing mEOS3.2:alpha-tubulin and SNAP:hCentrin2 was generated and selected using blastcidin and hygromycin. A related series of cell lines expressing various phosphomutant Hec1:SNAP under the control of a doxycycline promoter in addition to mEOS3.2:alpha-tubulin and SNAP:hcentrin2 were generated and selected with puromycin, blasticidin and hygromycin. An additional U2OS cell line with overexpressed mTurqouise2:Nuf2 and endogenously tagged beta-tubulin::tetracysteine(TC) as described previously (47) was utilized. A related series of cell lines expressing phosphomutant Hec1:LSS-mOrange under the control of a doxycycline promoter in addition to overexpressed mTurquoise2:Nuf2 and endogenously tagged beta-tubulin::TC were generated and selected using geneticin, blasticidin and hygromycin.

### Sample preparation and imaging conditions

Cells were plated onto coverglasses (GG-22-1.5-pdl, neuVitro) prior to experiments. Cells were preincubated in Fluorobrite DMEM (ThermoFisher) supplemented with 10mM HEPES for ∼15min immediately prior to imaging and then transferred to a custom-built coverslip heater maintained at 37°C during imaging. Cells were maintained in imaging media covered by mineral oil for ∼1 hour during experiments.

### Spinning Disk Confocal Imaging and Photoconversion

Photoconversion experiments were performed on a home built spinning disk confocal microscope (Nikon Ti2000, Yokogawa CSU-X1) with 488nm and 647nm lasers, an EMCCD camera (Hamamatsu) and a 60x oil immersion objective and controlled using a custom LabVIEW program described previously (71). Two fluorescence channels were acquired every 5s with either 500ms exposure, 561nm excitation, 593/40nm emission or 300ms exposure, 647nm excitation, 647 long pass emission in a single z plane for both channels. The mEOS3.2 was photoconverted using a 405nm diode laser aligned to a diffraction limited spot in the imagining plane of the microscope. The spot was moved across the sample at 5μm/s at a power of 500nW (measured at the objective) using a P-545 PInano XYZ; Physik Instrumente) to draw a photoconverted line across the spindle. Prior to experiments, cells were stained with 500nM SNAP-SiR (New England Biolabs) for 30 minutes and then recovered in culture media for at least 4 hours.

All quantitative analysis was performed using a custom MATLAB GUI described previously (12). We defined the spindle axis by tracking both poles and fitting a line between them. We then generated a line profile by summing over 3.5μm on either side of the spindle axis and fit the profile to a gaussian. The height of the peak was determined by subtracting the height of the fit gaussian from the height of a gaussian fit to the opposite side of the spindle to correct for background. The peak heights were then corrected for bleaching by dividing by a bleaching calibration curve. To measure the bleaching calibration curve, the entire spindle was photoconverted and then left to equilibrate for 5 minutes. The spindles were then imaged using the same imaging conditions as the photoconversion experiments. The mean intensity inside an ROI around the spindle was determined and averaged over 13 cells and subtracted the mean of a region outside of the cell to correct for the dark noise of the camera. We then divided the peak intensity vs time curve by the bleaching calibration curve to produce a bleaching-corrected intensity curve.

### Fluorescence Lifetime Imaging Microscopy – Fluorescence Resonance Energy Transfer (FLIM-FRET) Experiments

Fluorescence lifetime imaging microscopy (FLIM) experiments were performed on a home built two photon microscope as described previously (47). Prior to experiments cells were stained with 1μM FlAsH-EDT_2_ (ThermoFisher) in Opti-MEM for 2 hours, destained with 250μM EDT (Alfa Aesar) for 10 minutes, washed twice with Opti-MEM and then placed in culture media to recover for 6-10 hours. During imaging, cells were placed in a custom-built heating chamber and maintained at 37°C in imaging media for roughly one hour. Cells were continuously excited using an 80MHz Ti:Sapphire pulsed laser (Insight X3, MKS) tuned to 865nm using a galvanometer scanner (DCS-120, Becker & Hickl) with a 40x water immersion objective (CFI Apo 40x WI, NA 1.25, Nikon) for a total of 3 minutes of acquisition time split into 3s frames. FLIM signal was recorded using a TCPSC module (SPC-150, Becker & Hickl) with two hybrid detectors (HPM-100, Becker & Hickl). A dichroic beam splitter (FF506-Di03) split the emission signal between the two detectors. A 470/24nm bandpass emission filter was mounted on one detector and a 535/30nm bandpass filter on the other to record emission from mTurquoise2 and FlAsH respectively.

Quantitative FLIM analysis was performed using a custom MATLAB GUI described previously (47). Kinetochores were tracked in the mTurqouise2 channel using the Kilfoil tracking algorithm (97). The photons from each kinetochore pixel were then binned to a single FLIM decay curve for analysis and fit to a dual exponential using a Bayesian algorithm described previously (47, 98). The long lifetime of the dual exponential was fixed to 3.73ns, determined by fitting an mTurquoise2 FLIM decay curve from unstained cells to a single exponential. The fraction of NDC80c bound to KMTs was calculated by dividing the slow decay fraction from the fit dual exponential by a correction factor of 0.42 to account for the fraction of tubulin that was stained with the FlAsH FRET acceptor.

### Calculating the fraction bond f_bound_ for a cooperative two state system

We calculated the fraction of NDC80c bound to KMTs in a cooperative two-state system using the system’s partition function (99). We included 21 model states ranging from no NDC80c bound to all N=20 NDC80c bound to the KMT. We defined the energy of a state with *n* NDC80c bound, *U*(*n*), that included both a binding energy between NDC80c and KMTs, *ε*, and a pairwise, cooperative interaction energy between the NDC80cs bound KMTs, J: 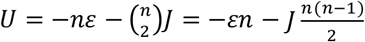. The partition function, Z, then becomes:

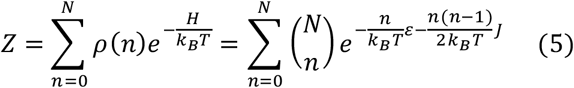

And

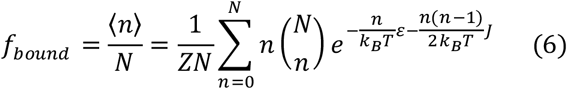

We defined an effective free energy difference between the states Δ*G* (66):

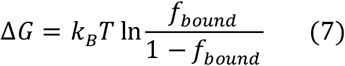

If there is no binding cooperativity (*J* = 0), the ratio of bound to unbound (i.e. the equilibrium constant) reduces to the familiar expression for a noncooperative two-state model:

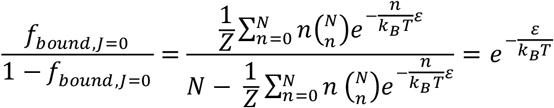

and the free energy difference between the two states becomes

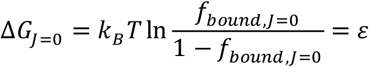

Furthermore, if the energy of binding varies linearly with the number of phosphorylated sites then *ε* = *ε*_0_ + *p*Δ*ε* where *p* is the number of phosphorylated sites, Δ*ε* is the change in binding energy per phosphorylated site, and *ε*_0_ is the binding energy of unphosphorylated NDC80c. Thus, if there is no cooperativity (i.e. *J* = 0) and if the energy of NDC80c-KMT binding varies linearly with the number of phosphorylated sites, then the free energy difference between bound and unbound NDC80c is predicted to also vary linearly with the number of phosphorylated sites (i.e. Δ*G*_*J*=0_ = *ε*_0_ + *p*Δ*ε*), as is observed experimentally (Figure 1F).

We also fit the experimentally observed free energy differences between the bound and unbound NDC80c states measured from FLIM-FRET experiments on the phosophomutants (Figure 1F) while allowing for the possibility of cooperativity (i.e. *J* ≠ 0). To do so, we again used the linear the model of NDC80c binding energy varying linearly with number of phosphorylated sites (i.e. *ε* = *ε*_0_ + *p*Δ*ε*) and calculated a predicted Δ*G* for each phosphomutants using Equations 6 and 7. We compared the predicted Δ*G* for each of the phosphomutants for a given *J*, Δ*ε, and ε*_0_ and fit these values to the free energy differences we measured in the FLIM-FRET experiment by minimizing a *χ*^2^ statistic. This resulted in best fit values of Δ*ε*=0.35 ± 0.03k_B_T and *ε*_0_=-0.50 ± 0.02k_B_T. The best fit value of NDC80-c-NDC80c interaction was *J*=-0.006 ± 0.004k_B_T, which was not significantly different from *J* = 0 (p=0.07 Bayesian likelihood ratio), indicating that NDC80c binding is noncooperative *in situ*.

### Expression of Hec1 phosphomutants with siRNA knockdown and doxycycline-induced mutant expression

U2OS cells were plated at 50% confluency in T75 flasks (ThermoFisher) in DMEM supplemented with 10% tetracycline-free FBS and PenStrep. We used a Flexitube siRNA duplex targeted to the 5’ UTR of the Hec1 gene (5’-TCCCTGGGTCGTGTCAGGAAA-3’, QIAGEN Hs_KNTC2_7 SI02653567) to knockdown endogenous Hec1. 240pmol of siRNA and 8μL of Lipofectamine RNAiMax (ThermoFisher 13778030) were separately incubated in two tubes of 1.2mL of Opti-MEM (ThermoFisher) for 5 minutes at room temperature and then combined and incubated for an additional 30 minutes of incubation. The DMEM media was replaced with 8mL of Opti-MEM supplemented with 10% tetracycline-free FBS. The siRNA-Lipofectamine cocktail was then added dropwise to the cell culture media. Cells were incubated with the siRNA media for 24 hours. We then washed the siRNA off and split the cells to six 35mm dishes containing 25mm poly-D-lysine coated coverslips and 2mL of DMEM supplemented with 10% tetracycline-free FBS supplemented with PenStrep and 2μg/mL of doxycycline to induce phosphomutant Hec1 expression for 24 hours before imaging. For experiments where additional genes were knocked down, a dual cocktail containing 240pmol of each siRNA and 16μL of Lipofectamine RNAiMAX incubated in 1.2mL of Opti-MEM was used. The following RNAis were used: MCAK: 50-50 mix of 5’-GCAGGCUAGCAGACAAAUUU and 5’-GAUCCAACGCAGUAAUGGUUU (Dharmacon); Kif18A: 5’-GCCAAUUCUUCGUAGUUUU (Dharmacon); Ska3: 5’-AACAGAATGGTTTAAGTAGTA (Qiagen).

### Materials Availability

All plasmids generated in this study are available on AddGene: 82793. All cell lines are available upon request from the Needleman Lab. DOIs are listed in the key resources table. Any additional information required to reanalyze the data reported in this paper is available from the lead contact upon request. All reagent and material requests should be referred to William Conway.

## DATA AVAILABILITY

All original code and data have been deposited at Dryad and are publicly available as of the date of publication

## ACKNOWLEDGEMENTS

This work was supported by the NSF-Simons Center for Mathematical and Statistical Analysis of Biology at Harvard (award number #1764269) and the Harvard Quantitative Biology Initiative. W. Conway was supported by National Science Foundation GRFP Fellowship # 2018238646. We thank Tae Yoon Yoo, Adam Lamson, and Maya Anjur-Dietrich for helpful comments on the manuscript.

---

**Supplemental Figure 1:**
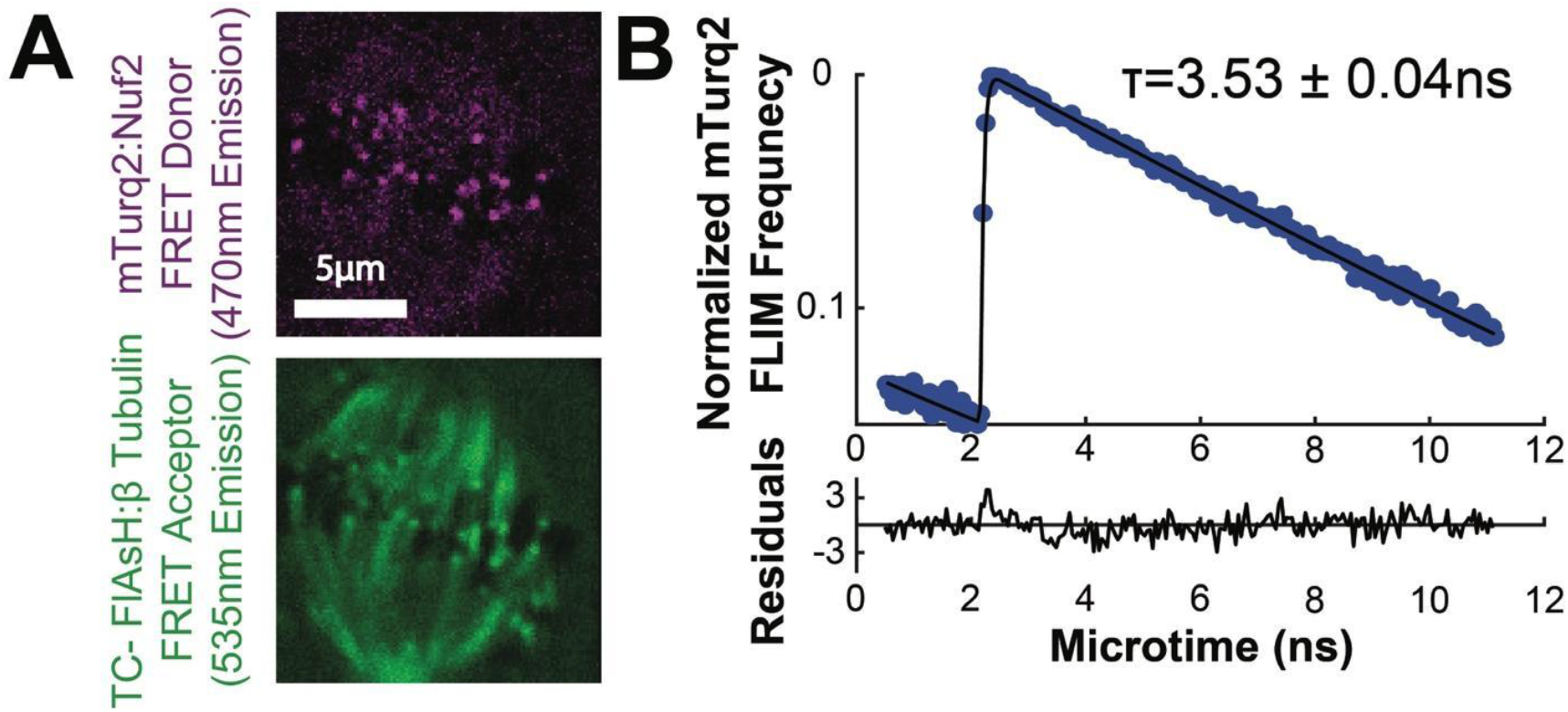
FLIM curve of the mTurquoise2 FRET donor in the presence of the FlAsH FRET Acceptor fit to a single exponential decay. A) mTurqouise2:Nuf2 FRET donor and the β-tubulin:TC-FlAsH FRET acceptor channels B) Normalized FLIM histogram of mTurquoise:Nuf2 localized at kinetochores fit a single-exponential

**Supplemental Figure 2:**
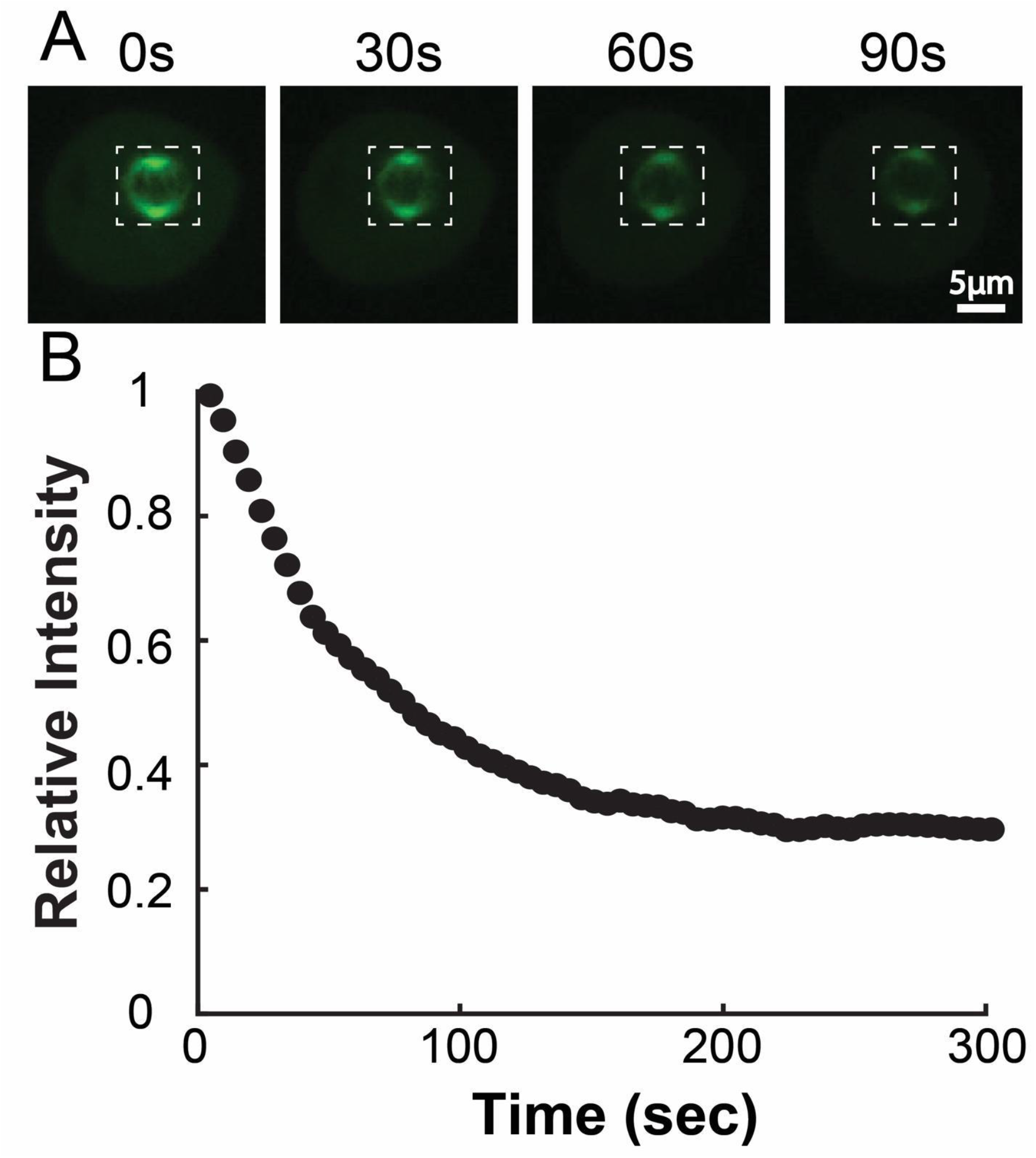
mEOS3.2: *α*-tubulin bleaching correction curve. A) Time series of activated tubulin in spindles. The whole spindle was photoactivated with a 405nm laser and left to equilibrate for 5 minutes before imaging B) Mean integrated spindle intensity over time (boxed region). Curves were corrected for dark background by subtracting the mean intensity of a small region marked outside the cell. Curves from 13 cells were normalized to the initial intensity at t=0s and then averaged together.

**Supplemental Figure 3:**
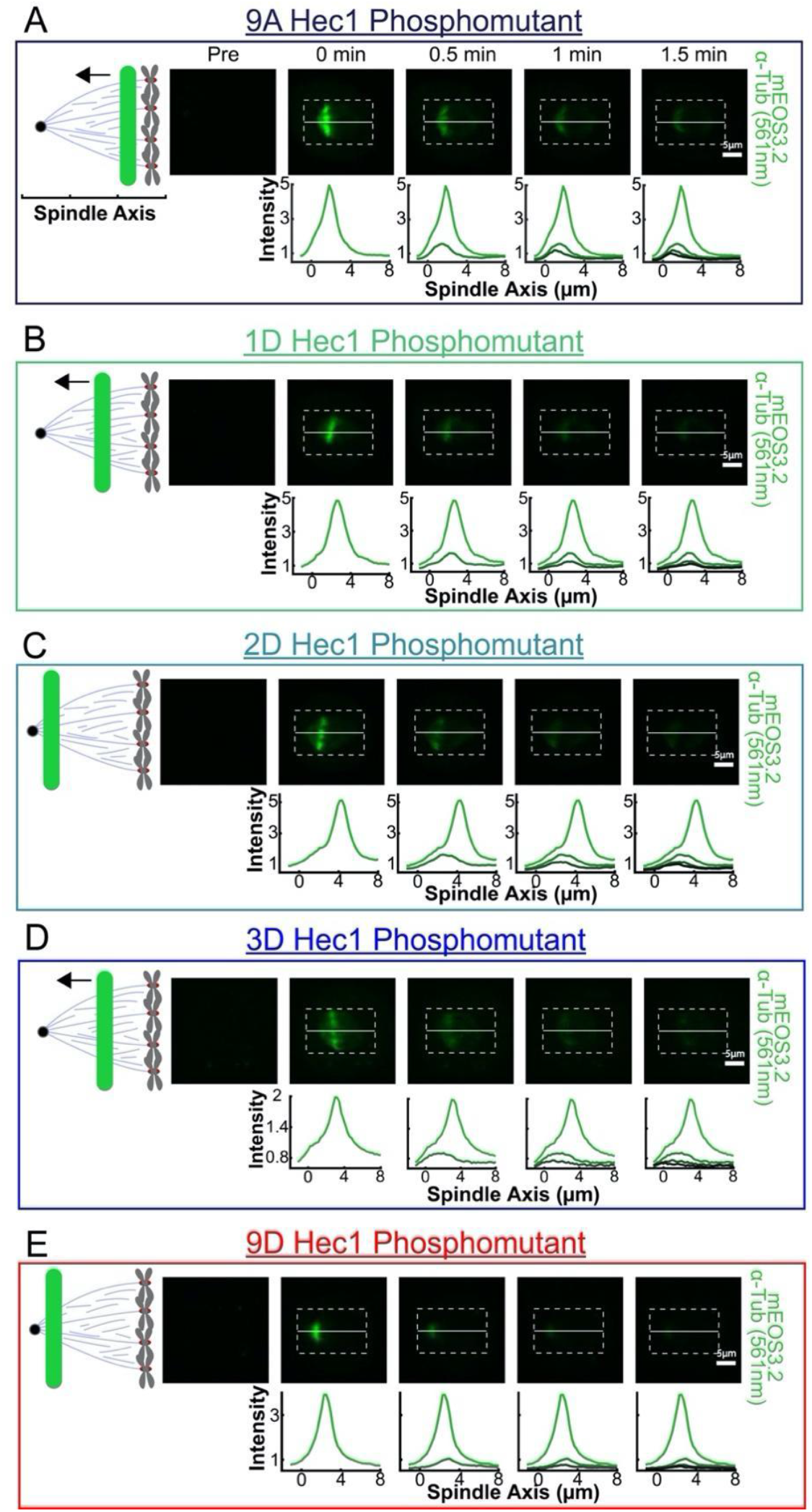
Photoactivation line profiles for Hec1 Phosphomutants. **A**) Images from a photoconversion experiment in cells expressing a 9A-Hec1 Phosphomutant showing SNAP: mEOS3.2:α-tubulin (green). Images are shown immediately preceding, 0, 30, and 60s after photoconversion with a focused 405nm diode laser Line profile generated by averaging 3.5μm on either side of the spindle axis in the dotted box. B) 1D-Hec1 C) 2D-Hec1 D) 3D-Hec1 E) 9D-Hec1

**Supplemental Figure 4:**
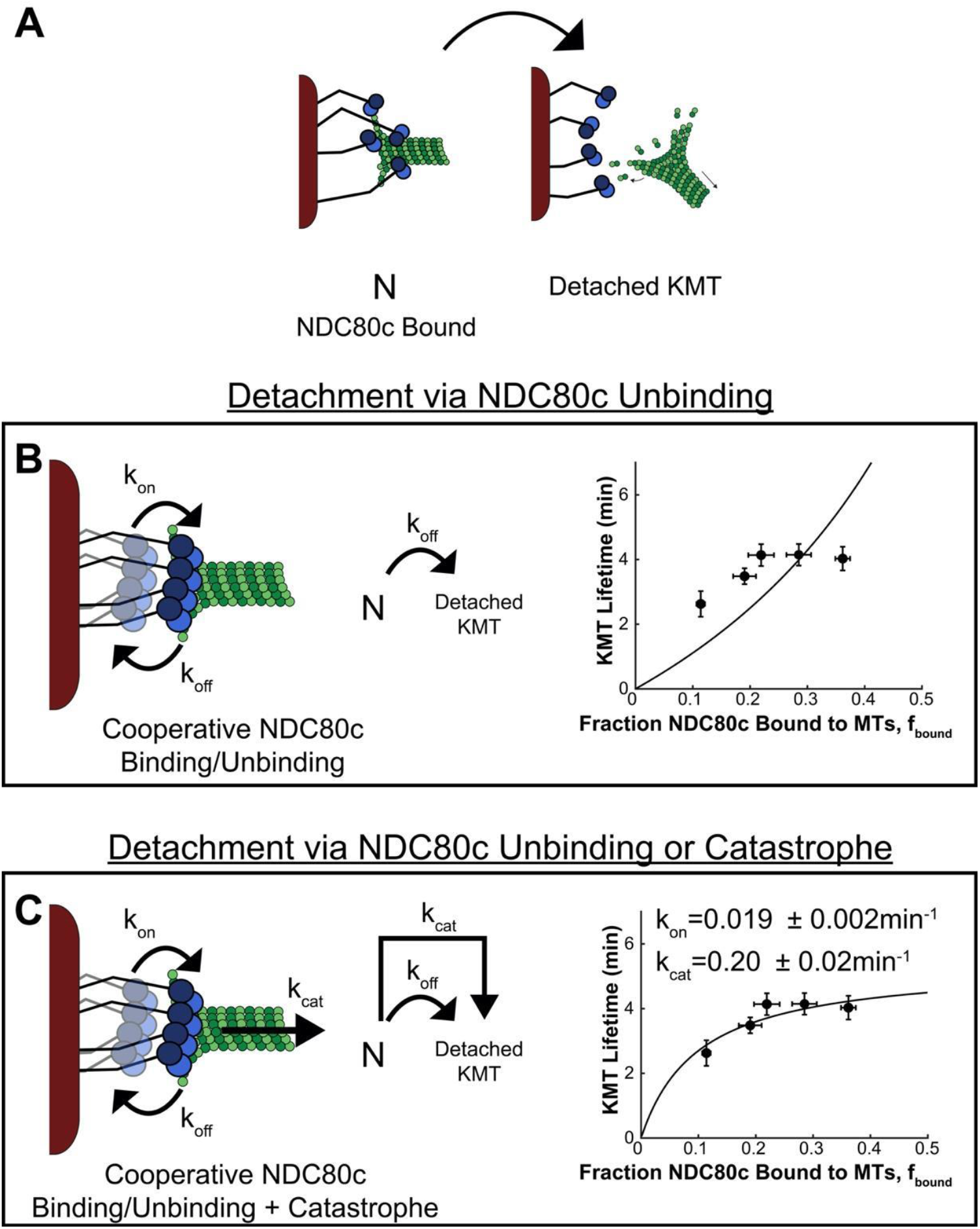
KMT detachment models with cooperative NDC80c binding. A) A rate transition model with all NDC80c bound and a KMT detached state B) Left: A model with cooperative NDC80c binding and unbinding. Middle: Transitions between states defined only by NDC80c binding *k*_*off*_. Right: KMT Lifetime vs Fraction of NDC80c bound to MTs for Hec1 phosphomutants (black dots) compared with model predictions (black line). C) Left: A model with cooperative NDC80c binding and unbinding and KMT catastrophes. Middle: Transitions between states defined by NDC80c binding: *k*_*off*_ and by KMT catastrophe: *k*_*cat*_. Right: KMT Lifetime vs Fraction of NDC80c bound to MTs for Hec1 phosphomutants (black dots) compared with model predictions (black line)

## Appendix 1: Calculation of the mean first passage time in the KMT detachment model

Here, we derive the analytic expressions for the mean KMT lifetime (Equations 2 and 3 in the main text) and compare these analytic expressions with full numerical calculations. We consider a model where there is a fixed number, *N* = 20, of NDC80cs per KMT. Each NDC80c is either bound or unbound to the KMT. Each NDC80c independently binds and unbinds to/from the KMT at rates *k*_on_ and *k*_off_. The KMT detaches when either all the NDC80cs unbind from the KMT or following catastrophic detachment happening at a constant rate *k*_cat_ that is independent of NDC80c binding. Since the kinetochores are in steady state, new KMTs must nucleate to replace the detaching KMTs. The nucleation process will be discussed in detail in Section III. An illustration of the model is shown in Figure A1.

Since there are *N* copies of the NDC80c per KMT, we treat the system as a Markov chain with *N* + 1 states of 0, 1, 2, …, *N* NDC80c bound. The case *k*_on_ = *k*_off_ is considered in the classic problem of Ehrenfest urn [1–3]. In the Ehrenfest urn problem, *N* particles independently move between two containers at a fixed rate. All *N* particles initially begin in one of the containers and the expected time to return to the initial state of all *N* particles in one container is calculated. The more general case with *k*_on_ ≠ *k*_off_ has also been previously considered [4–6] and a generalized formula has been derived in Ref [6]. For completeness, we provide here an alternative, self-contained derivation. We first discuss the case without catastrophic events, leading to the useful expression of Equation 21. We then extend the results to the case with catastrophic events and provide a simplified approximate expression in the regime relevant for the kinetochore experiments. Finally, we compare these approximate expressions with numerical solutions and experimental results.

**Fig. A1:**
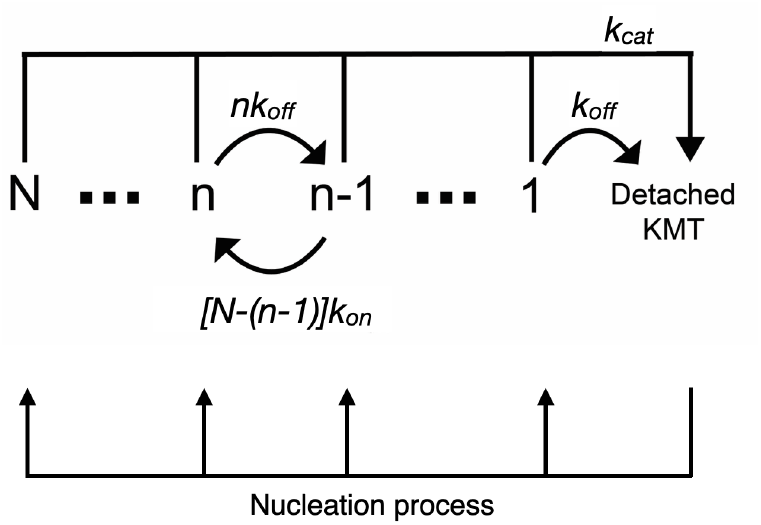
Schematic of the full detachment-attachment model.

### I. CALCULATION OF THE MEAN KMT LIFETIME IN THE DETACHMENT ONLY VIA NDC80C UNBINDING MODEL

**Fig. A2:**
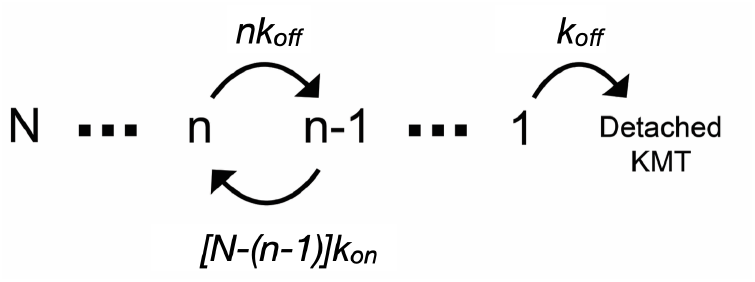
Schematic of the detachment only via NDC80c unbinding model.

We first consider a model where KMTs only detach when all NDC80c simultaneously unbind (i.e. the catastrophe rate is set to zero, Figure A2). For simplicity, we non-dimensionalize to set the unit time as *t*_0_ = 1*/k*_off_. The forward rate (the rate for each NDC80c to go from bound state to unbound state) is then 1 and backward rate is *b* = *k*_on_*/k*_off_. We define *P*_*n*_ as the probability of being in the *n*^th^ state with *n* NDC80c bound. Since the *n*^th^ state contains *n* NDC80c, which can each independently unbind, the probability of transitioning from the *n*^*th*^ state to the (*n* −1)^*th*^ state in a unit time step *t*_0_ is *n*, where the *k*_off_ unbind rate has been absorbed by the non-dimensional time unit *t*_0_. The probability of transitioning from the *n*^*th*^ state to the (*n* + 1)^*th*^ state is *b*(*N* − *n*). The transition matrix *T* = {*T*_*n,m*_}, where each entry *T*_*n,m*_ represents the probabilities of moving from one state *n* to another state *m*, takes the form:

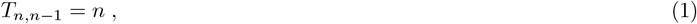

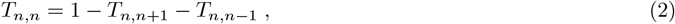

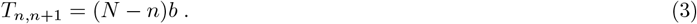

Equivalently, the forward equation describing how the probability distribution over different sites *P*_*n*_ changes with time is:

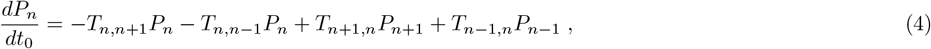

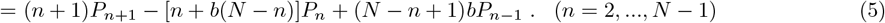

We define *x*_*n*_ as the mean first passage time for the system to go from the *n*^th^ state (with *n* NDC80c bound) to the 0^th^ state (detachment). Using a recurrence relation, we can see that 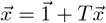. With the transition rates *T*_*n,m*_ given above, this leads to:

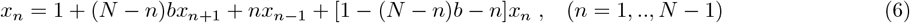

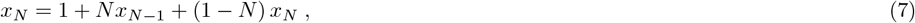

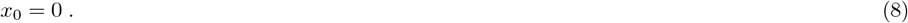

Next, we define *z*_*n*_ = *x*_*N*−*n*+1_ − *x*_*N*−*n*_ (*n* = 1, ‥, *N*) to get the following recurrence relation:

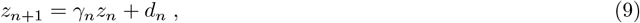

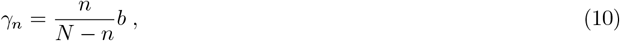

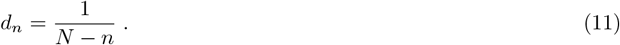

The initial condition is 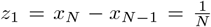, which can be seen from Equation 7. The mean first passage time for the system to go from the (*N* − *n*)^th^ state (with *N* − *n* NDC80c bound, *n* NDC80c unbound) to the 0^th^ state (detachment) is 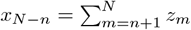.

It is helpful to define the auxiliary variables:

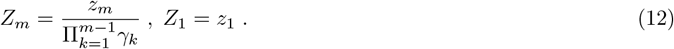

Thus we have

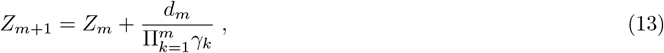

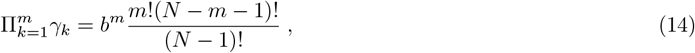

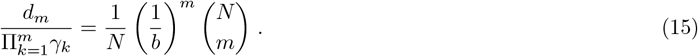

So the expression for *Z*_*m*_ is

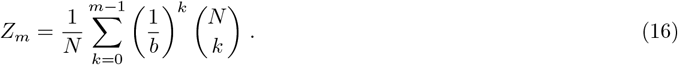

Here we define 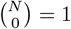 for convenience. Then we have,

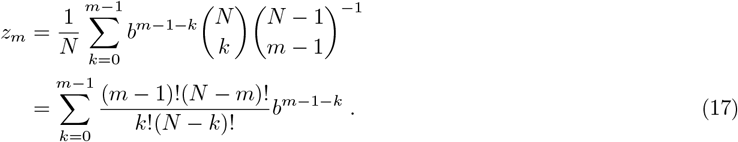

Therefore we have the mean first passage time for the system to go from (*N* − *n*)^th^ state (where *N* − *n* units are bound, *n* units are unbound) to 0^th^ state (detachment):

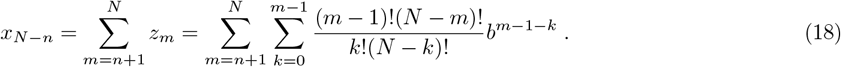

In particular, we have the expression for *x*_*N*_

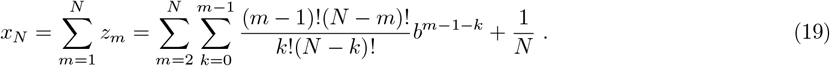

For a system with *N · b* ≫ 1, the first passage time for the system to go from the all-bound state to the non-bound state is dominated by the last step where the final NDC80c unbinds. This approximation can be inferred from Equations 9 and 10 and we will further verify this approximation numerically. This means that, in fact, the mean first passage time is largely independent of the initial condition. In this case, the distribution of first passage times will be exponential with a mean of *x*_1_, as the first passage time distribution is dominated by repeated, comparatively slow, independent attempts to detach the final NDC80c. The mean time for the system to transition from the final 1 NDC80c bound state to the detached state, *x*_1_, which is a good approximation for the mean lifetime of the KMT when *N* · *b* ≫ 1, is:

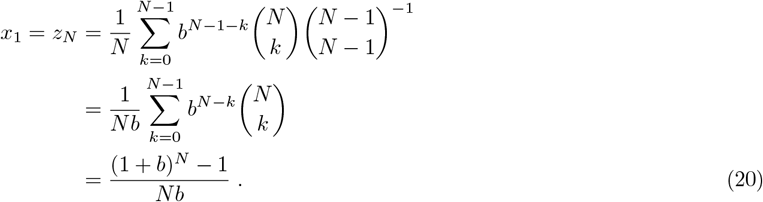

Putting back the dimensions, we obtain the following.

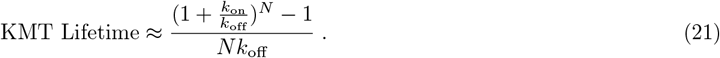

### II. CALCULATION OF THE MEAN KMT LIFETIME IN THE DETACHMENT VIA NDC80C UNBINDING OR FOLLOWING KMT CATASTROPHE MODEL

Now we consider a model where the KMT detaches when either the final NDC80c unbinds from the KMT or following a KMT catastrophe. When a catastrophic event occurs, the system transitions directly from a state with *n* NDC80c to the detached state without transitioning through the intermediate *n* − 1, *n* − 2, …, 1 states, at a rate *k*_cat_ (Figure A3). This adds a *k*_cat_ term to each of the forward equations for transitions between states.

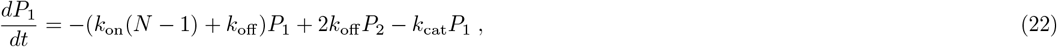

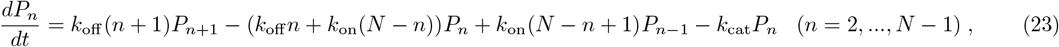

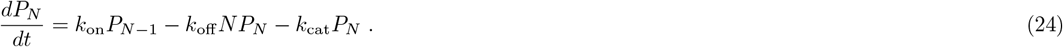

**Fig. A3:**
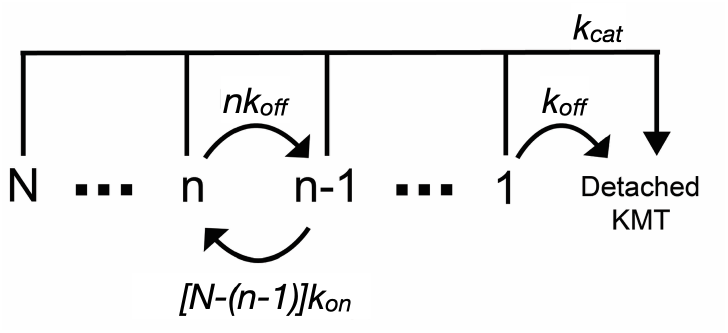
Schematic of the detachment via NDC80c unbinding or KMT catastrophe model.

We define a change of variable 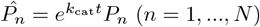 and observe that 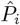 denotes the transition probabilities of a similar system but without the catastrophe rate. To see this, we re-express the above equations as 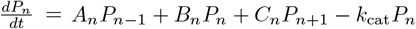. Then we have

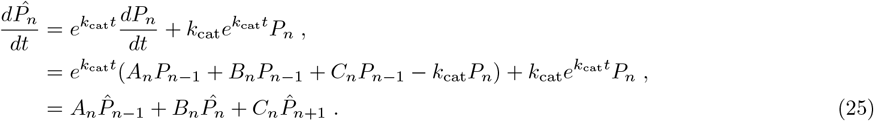

This change of the variable 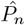 thus transforms the system back to the original equations without the catastrophic events. The net effect of the catastrophic events is to decrease the probability of staying in each state with *n* NDC80c bound (*n* = 1, …, *N*) by a factor of 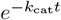, implying that 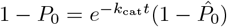. (Here 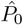 denotes the probability to arrive at the detached state 0 with the same initial condition in a system without catastrophic events.) For the system with catastrophes, the first passage time distribution can be approximated by adding the two independent NDC80c unbinding and catastrophe rates, yielding an exponential distribution with mean lifetime:

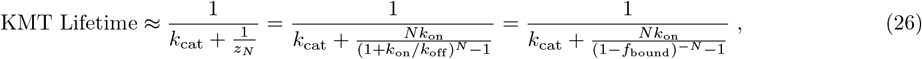

where the final equality comes from the relation: *f*_bound_ = *k*_on_*/*(*k*_on_ + *k*_off_). With catastrophe driven detachment included, the first passage time is even less sensitive to the initial condition, which justifies the first approximate equality in the equation above. We fit the *k*_cat_ and *k*_on_ values in this equation to the FLIM-FRET and KMT lifetime data and found that *k*_on_ = 0.08 min^−1^ and *k*_cat_ = 0.25 min^−1^ (see Figure 2G, main text). Intuitively, we can extract the *k*_cat_ value from the height of the KMT lifetime plateau as *f*_bound_ increases, since the second term in the denominator of the rate approaches zero as *f*_bound_ grows. We can extract the *k*_on_ value by noting that the KMT lifetime is half of its maximum when the two terms in the denominator are equal, implying that *k*_on_ = *k*_cat_((1 − *f*_bound_)^−*N*^− 1)*/N* at the half-maximum point. This means that *k*_on_ value can be estimated from the *k*_cat_ value (itself estimated from the lifetime plateau at high *f*_bound_), the number of NDC80c, *N*, and the value of *f*_bound_ where the KMT lifetime reaches its half-maximum.

### III. NUMERICAL CALCULATION OF THE DETACHMENT PROCESS

Here we calculate the KMT lifetime numerically to verify the results we derived in the previous sections. To do so, we must specify the initial condition of the system. The number of KMTs is very similar throughout mitosis [7] and the lifetime of an individual metaphase KMT (∼ 4 min) is much less than the duration of mitsosis in human cells (∼ 45 min). This implies that the system is in a steady state during metaphase. That is, statistical properties such as the distribution of the number of NDC80 complexes bound to each kinetochore should remain consistent and time-independent, regardless of when we begin to measure KMT lifetime in our experiments. We therefore take the steady state distribution of the number of bound NDC80cs as the initial condition used to calculate the KMT lifetime, which will be further detailed in Equation 28-30. In steady state, the number of KMTs is constant, so new KMTs must nucleate to replace the detaching KMTs. We assume that each kinetochore has a fixed number of sites *N*_KMT_ where KMTs can bind. These sites could be occupied by a KMT or unoccupied, with a mean number of KMTs at the kinetochore *n*_KMT_. In steady state, the nucleation rate *k*_nucl_ and the detachment rate *k*_detach_ are related by: *n*_KMT_*k*_detach_ = (*N*_KMT_ − *n*_KMT_)*k*_nucl_. In prior experiments, stabilizing KMTs by treatment with taxol, a microtubule-stabilizing drug which inhibits microtubule catastrophes, only slightly increases the number of KMTs in metaphase [7]. This implies that the kinetochore is nearly saturated (i.e. *N*_KMT_ ≈ *n*_KMT_) and that the nucleation rate is much larger than the KMT detachment rate *k*_nucl_ ≫ *k*_detach_. Since the detachment rate is the inverse of the mean KMT lifetime (*k*_detach_ = 1*/*(KMT Lifetime) ≈ *k*_cat_), we set *k*_nucl_ = 3min^−1^ ≈ 10*k*_cat_. We will systematically examine the impact of varying the *k*_nucl_ parameter on the NDC80c fraction bound and steady state distributions after solving the system numerically for a given value of *k*_nucl_.

Since the kinetochore is a large complex structure, the NDC80c binding and unbinding from the KMT must involve large-scale rearrangements of many subunits within the kinetochore. Fluctuations between these two states should continue regardless of whether the site is occupied by a KMT or not, implying that each NDC80c at an unoccupied site should continue to transition between a “bound” state where it is located near KMT, and therefore be available to immediately bind to a freshly nucleated KMT, or an “unbound” state where it is located far from the KMT and not available to immediately bind to the KMT. For simplicity, we assume that the transition rates between the two states will be the same regardless of whether the KMT is present or not, though the interaction between the NDC80c and the KMT could certainly alter the energy landscape of the NDC80c. Since each NDC80c will continue to independently transition between the “bound” and “unbound” states even when the KMT is absent, the rate that the KMT nucleates into the *n*^th^ state with *n* NDC80c bound will be given by a binomial distribution set by the on and off rates of the NDC80c binding:

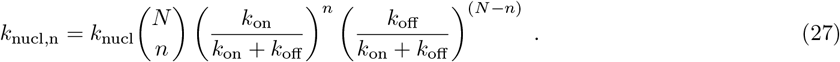

Then we can write down the equations for the steady state probability distribution *P*_eq,n_ including the nucleation terms (where *P*_eq,n_ is the probability to be in state *n* with *n* NDC80c bound):

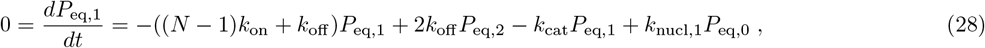

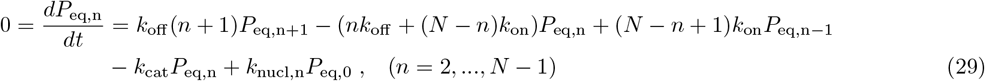

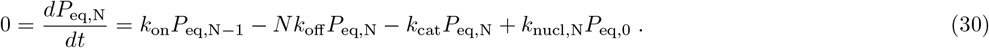

And *P*_eq,0_ is given by 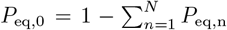. We numerically solved these linear equations to find the steady state distribution of NDC80c bound (Matlab linsolve function, which uses LU factorization with partial pivoting - full script available in https://github.com/LuyiQiu/KMT_lifetime). We then used this steady state distribution as the initial condition and numerically calculated the first passage time probability distribution for the system to get to detached state (i.e. the KMT lifetime) using the eigenvalue method (- full script available in https://github.com/LuyiQiu/KMT_lifetime). The numerical results show that the lifetime distribution is well approximated by an exponential with a mean given by the only-last-step analytic solution (Equation 26) at the best fit parameter values (Figure A4, *k*_cat_ = 0.25min^−1^, *k*_on_ = 0.08min^−1^).

As a final consistency check and to set a bound on the value of *k*_nucl_, we consider another taxol experiment in ref [8]. This work shows that treating the 9A-Hec1 mutant with taxol did not significantly alter the fraction of NDC80c bound to KMTs (less than 8% change within error, Figure 5A in Ref [8]), implying that setting *k*_cat_ = 0 will not increase *f*_bound_ for the 9A-Hec1 phosphomutant by more than 8%. To test this, we computed the relative change of fraction of NDC80c bound as a function of the nucleation rate *k*_nucl_ when we switch *k*_cat_ from 0.25 min^−1^ to 0. The restriction that the fraction of NDC80c increased by less than 8% requires *k*_nucl_ *>* 3 min^−1^ ≈ 10*k*_cat_(Figure A5A). Indeed, further increase *k*_nucl_ would barely change the outcomes. This confirms that within our current assumption of the nucleation process, the results are insensitive to the choice of *k*_nucl_. We also plot the steady state distribution of NDC80c bound to KMTs at *k*_nucl_ = 3 min^−1^ (Figure A5B) for both the strongest and weakest binding mutants (9A-Hec1 and 9D-Hec1). We found that in both cases the steady state distribution is well approximated by binomial, which verifies the equality we used 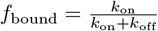 in deriving Equation 26.

We then numerically calculated the relationship between the KMT lifetime and fraction of NDC80c bound at the best fit parameters (Figure A6). The analytic, numerical and experimental curves are in good agreement, indicating that a model where KMTs detach from the kinetochore when all NDC80c simultaneously unbind or following a KMT catastrophe is consistent with the fraction of NDC80c and the KMT lifetime and that the only-last-step approximation is reasonable for the range of NDC80c fraction bound measured in the experiments.

**Fig. A4:**
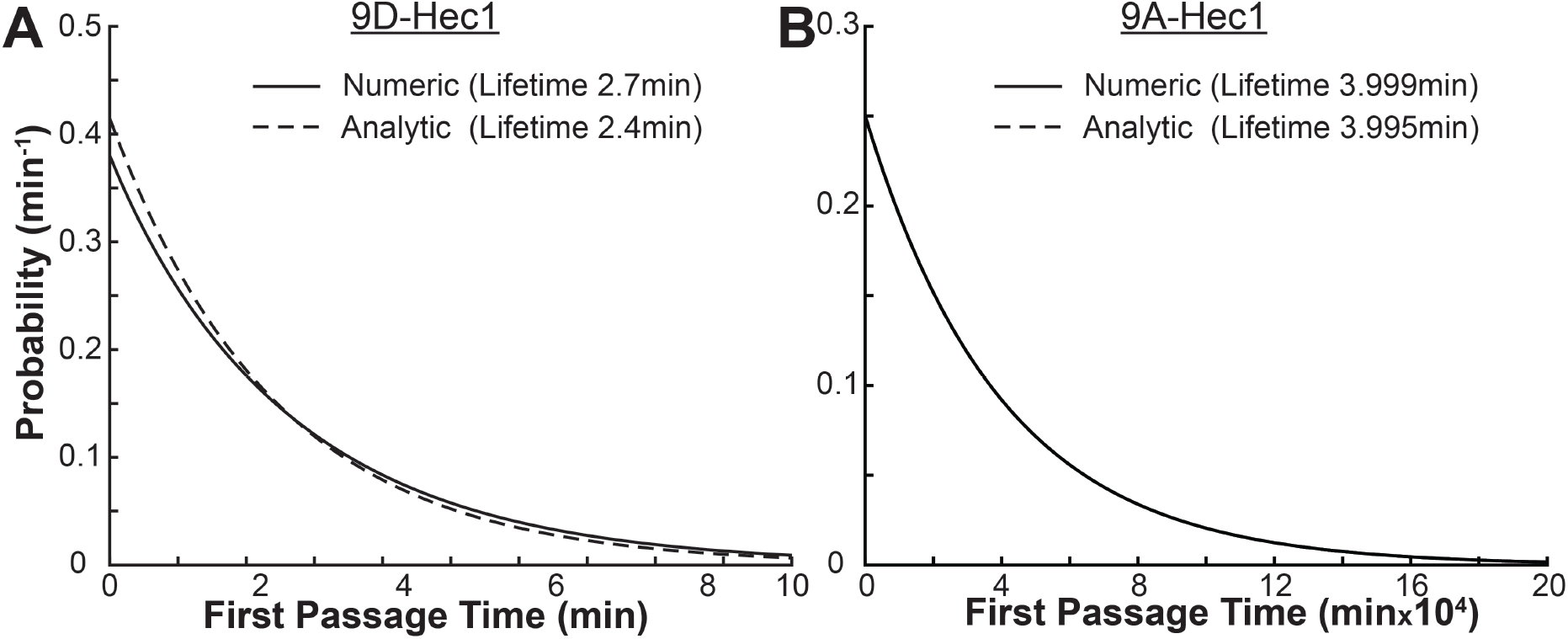
Distribution of first passage detachment times for the NDC80c unbinding or catastrophe model A) Comparison of full numeric and only-last-step approximate analytic solution for the first passage detachment time for the 9D-Hec1 mutant (*Nk*_on_*/k*_off_ = 2.4). The only-last-step analytic solution prediction (2.4min) is 10% lower than the exact numeric prediction considering all transition steps (2.7min). B) Comparison of full numeric and only-last-step approximate analytic solution for the first passage detachment time for the 9A-Hec1 mutant (*Nk*_on_*/k*_off_ = 13.2). The only-last-step analytic solution prediction (3.995min) is 0.1% lower than the exact numeric prediction considering all transition steps (3.999min).

**Fig. A5:**
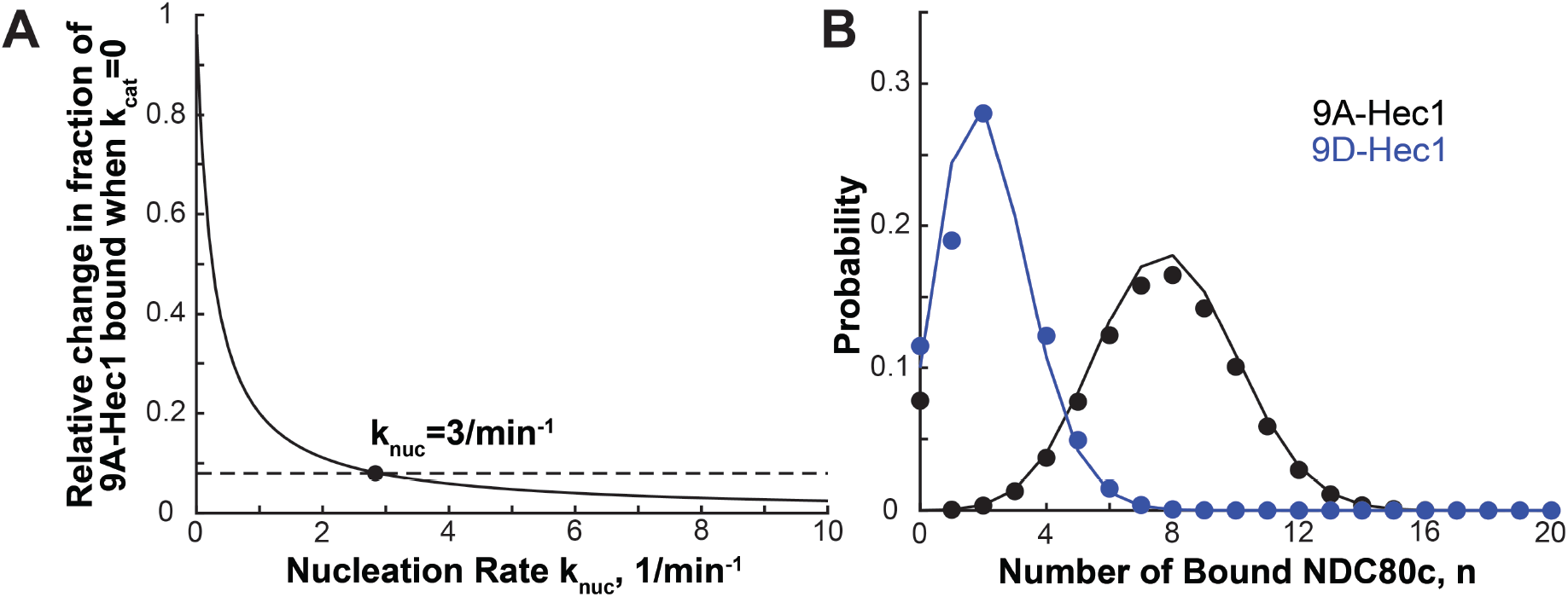
A) The change in the fraction of 9A-Hec1 NDC80c bound to KMTs when changing *k*_cat_ from 0.25min^−1^ to 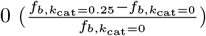 vs. the nucleation rate *k*_cat_. Dashed line: Maximum change observed in FLIM-FRET experiments in Ref [8]. The change in the fraction of 9A-Hec1 NDC80c is less than the cutoff when *k*_nuc_ *>* 3min^−1^ B) Comparison of numerically calculated steady state distribution of number of NDC80c bound to KMTs at the *k*_nucl_ = 3 cutoff value computed numerically for 9D-Hec1 (blue) and 9A-Hec1 (black) phosphomutant NDC80c. Blue: 9A-Hec1 with binomial distribution. Dots: Numerical calculation; Line: Binomial distribution.

**Fig. A6:**
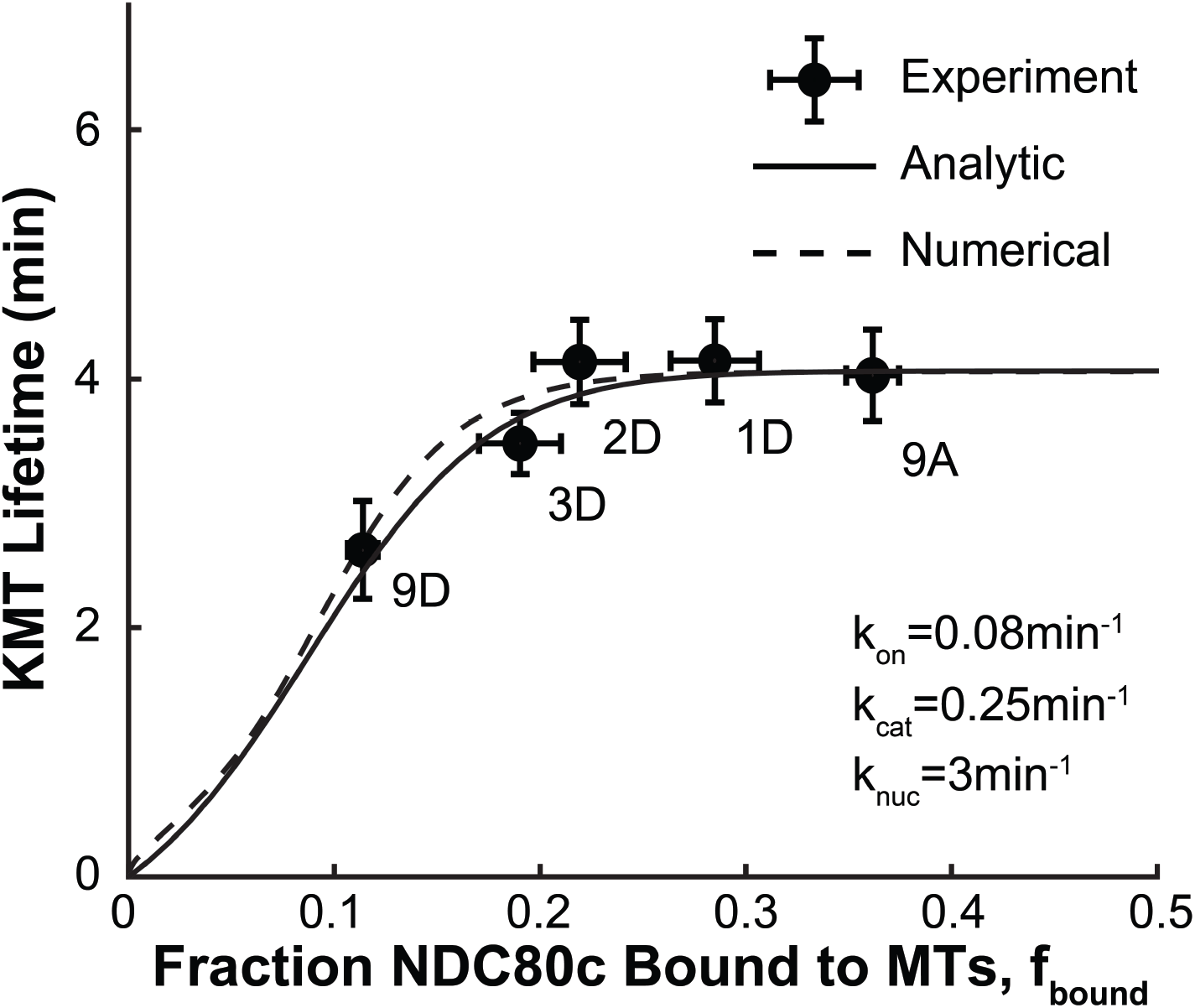
Comparison of the only-last-step approximation (Equation 26) and numerical calculation of the KMT lifetime vs faction of NDC80c bound curve in the detachment via NDC80c unbinding or following KMT catastrophe model. Dots: Experimental measurements; Solid Line: Only-last-step analytic approximation (Equation 26); Dashed Line: Numerical calculation

